# Hippocampal dorsal and ventral CA2 enable social discrimination after acute social defeat

**DOI:** 10.64898/2026.11.09.765487

**Authors:** Wanhui Sheng, Lara M. Boyle, Domenico Pimpinella, Pegah Kassraian, Haoyuan Li, Phoebe Matthew, Steven A. Siegelbaum

## Abstract

Remembering the valence of conspecifics based on past aggressive compared to non-aggressive interactions is important for proper adaptive social behaviors. How a single naturalistic encounter establishes a lasting memory of the valence of the experience remains unclear. Our recent work shows that the CA2 region in ventral hippocampus, unlike dorsal CA2, is not required for social novelty discrimination but does contribute with dorsal CA2 to promote social aggression. How CA2 contributes to valence-associated social memory across the dorsoventral axis remains unresolved. We developed an acute defeat-associated social threat discrimination memory (AD-STDM) paradigm after which a male mouse preferentially interacted with a familiar non-aggressor male compared to a familiar aggressor male 24 h after a single defeat episode. Chemogenetic inhibition of Avpr1b-expressing CA2 neurons in either dorsal or ventral hippocampus during learning impaired subsequent discrimination. Acute defeat also increased cFos expression in ventral CA2 neurons relative to non-aggressive social interaction. Thus, although ventral CA2 is not required for social novelty recognition memory, it is required for social memory of an aversive social experience, with a relatively greater level of neural activation during aversive compared to non-aversive social encounters. In contrast, dorsal CA2 is required for both non-aversive and aversive forms of social memory.

## Introduction

Adaptive social behaviors depend on the ability of an animal to recognize conspecifics and remember the outcomes of previous encounters. Non-aggressive social interactions can be rewarding, whereas aggressive encounters can lead to subsequent avoidance^1,2^. Memories linking individuals to these experiences provide a basis for distinguishing conspecifics associated with different social emotions and guiding subsequent approach and avoidance. Failure to build a correct social perception can lead to unnecessary conflicts and disrupted social behavior in many psychiatric disorders^3–5^. Understanding how the brain forms and retrieves such memories is therefore essential for explaining how social experience shapes behavior in healthy subjects and the neuronal bases for how this process may be disrupted in pathological conditions.

For more than a decade, dorsal hippocampal CA2 (dCA2) pyramidal neurons have been recognized as being important for social novelty recognition memory (SNRM) in rodents^6–8^, the ability of an animal to discriminate a novel from familiar conspecific. Much less is known about the function of CA2 in ventral hippocampus (vCA2). Indeed, whether a distinct CA2 region extends into the ventral hippocampus was controversial because of the inconsistent expression of dCA2 markers across the dorsoventral axis^9–11^. Our recent work provided multiple lines of evidence for the existence of a distinct CA2 region that extended throughout ventral hippocampus. Ventral CA2 shares certain molecular, electrophysiological, anatomical and behavioral properties with dCA2, but also shows some fundamental differences in gene expression, synaptic connectivity, and behavioral function^12^. This mirrors other studies showing that dorsal and ventral CA1 (dCA1 and vCA1) regions in hippocampus have differences in molecular and behavioral functions^13^. At a behavioral level, vCA2 pyramidal neurons were not required for SNRM. However, both dCA2 and vCA2 promote social aggression, an emotionally driven social behavior, with dCA2 acting through its projection to the lateral septum^12,14^. This difference is consistent with the known distinction along the hippocampal dorsoventral axis: Whereas dorsal hippocampus is more important for cognitive memory^15,16^, ventral hippocampus is more important for emotional behaviors, including aversive fear memory and anxiety-like behaviors^17–21^. These functional distinctions have been well established in CA1 and CA3, but the function of CA2 along the dorsoventral axis has been far less studied. Here we investigate the hypothesis that vCA2 is more important for social discrimination memory of naturalistic threat-associated social encounters.

Recent studies implicate dCA2 in social memory and behavior associated with aversive experiences. In a foot-shock-paired social fear-conditioning paradigm, dCA2 activity was required during both learning and recall for discrimination between threat-associated and safety-associated conspecifics^22^. Dorsal CA2 also contributes to behavioral responses following acute social defeat (AD): inhibiting dCA2 pyramidal neurons during defeat diminished a short-term memory of the encounter, assessed by the preferred interaction of the subject towards a familiar non-aggressor conspecific compared to the aggressor when tested 5-min after the encounter^23^. However, it is unclear from this study whether a subject can encode a longer-term social threat discrimination memory of the aversive compared to non-aversive social experience.

Consistent with the involvement of the ventral hippocampus in emotionally relevant behaviors, ventral CA1 (vCA1) has been recently found to be important for social threat discrimination memory associated with acute social defeat. Zorab and colleagues found that serotonergic inputs from the dorsal raphe and neurotensin inputs from the paraventricular thalamus to vCA1 support positive and negative social valence memory, respectively^23^. Suto and colleagues subsequently demonstrated that updating the negative valence of a familiar individual involves strengthened connections between vCA1 social-memory neurons and basolateral amygdala neurons carrying negative valence, within a circuit involving the nucleus accumbens^24^. These studies establish mechanisms through which vCA1 supports experience-dependent social valence, while leaving the contribution of vCA2 unresolved. Together, these findings motivate our current investigation into how dorsal and ventral CA2 contribute to discrimination between individuals previously encountered during aggressive and non-aggressive interactions.

## Results

### Acute defeat-associated social threat discrimination memory (AD-STDM) test

To investigate how natural social experiences of different valences lead to long-term social threat discrimination memory, we developed an acute defeat-associated social threat discrimination memory (AD-STDM) test (Fig. 1a test). During a Day 1 learning session, a male subject mouse (C57BL/6J) interacted with a novel non-aggressive (NA) male BALB/c stimulus mouse in a 10 min social interaction session in a neutral arena; the subject was then returned to its home cage. After 30-minutes, the subject was introduced into the home cage of a novel aggressive (A) male BALB/c mouse using a standard resident-intruder test, leading to the acute defeat (AD) of the subject (intruder) by the resident (aggressor) over a 10 min session. In this way the subject underwent two social experiences of opposing valence (moderately positive versus aversive; Supplementary Fig. 1). To maximize chances for the desired non-aggressive or aggressive social encounters, the A mice were prescreened for their tendency to engage in the appropriate behaviors during interactions with a separate group of C57BL/6J males. During a Day 2 recall session, 24 h later, the subject explored the now familiar NA and A stimulus mice confined to wire cup cages placed in opposite side chambers of a three-chamber arena. The subject mice spent significantly more time interacting with the NA mouse than with the A mouse (Fig. 1b), with a social discrimination index (SDI), (time exploring NA – time exploring A)/(total exploration time), significantly greater that zero (one-sample t test: t = 4.08, p = 0.0009; Fig. 1c).

**Figure 1.**
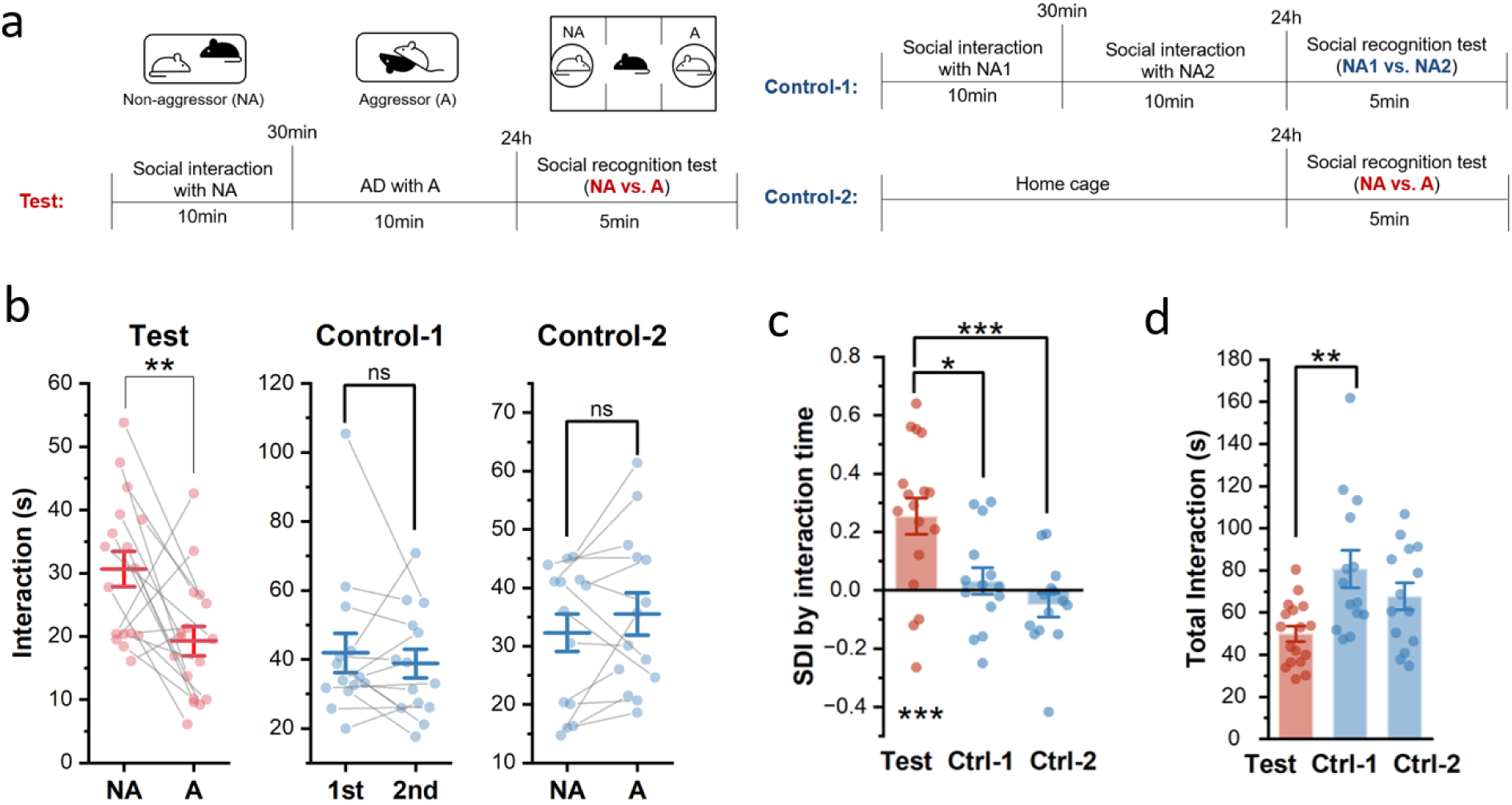
AD leads to long-term social discrimination memory. **a.** Schematic of the AD-STDM test and control conditions. During the day 1 learning session, test mice underwent a 10-min social interaction with a NA BALB/c, followed 30 min later by a 10-min AD by an A BALB/c. 24 hours later, each subject mouse underwent a 5-min social-recognition test in a three-chamber arena, in which the familiar NA and A were confined beneath wire cups in opposite side chambers. In control-1, subject mice interacted sequentially with two different non-aggressors (NA1 and NA2), separated by 30 min, without undergoing AD. 24 hours later, social recognition was assessed using NA1 and NA2. control-2 mice remained in their home cage on day 1 and underwent the social-recognition test with a novel pair of NA and A 24 h later. **b.** Direct interaction time with the social stimuli. Test mice spent more time interacting with the familiar NA than with the A (left). Control-1 mice showed no difference in interaction time between the first and second NA, indicating that presentation order did not produce a preference (middle). Control-2 mice showed no difference in interaction time between the NA and A in the absence of day 1 learning (right). (n = 17 in test, 14 in ctrl-1, 14 in ctrl-2; paired t test, test: t = 3.16, p = 0.0061; ctrl-1: t = 0.66, p = 0.5208; ctrl-2: t = −1.249, p = 0.234) **c.** Social discrimination index (SDI) was higher in the test group than in control-1 and control-2. (n = 17 in test, 14 in ctrl-1, 14 in ctrl-2; One-way ANOVA followed by Tukey’s multiple-comparisons test for across-group comparison; one-sample t test for comparison vs. 0, F(2, 42) = 9.512, p = 0.0004, test vs. ctrl-1 p = 0.01, test vs. ctrl-2 p = 0.0004; t(test) = 4.08, p = 0.0009.) **d.** Total direct interaction time during the social-recognition test. Test mice exhibited a lower total interaction time than control-1 (One-way ANOVA followed by Tukey’s multiple-comparisons test, F(2, 42) = 6.173, p = 0.0045, test vs. ctrl-1 p = 0.003). *p < 0.05, **p < 0.01, ***p < 0.001

Because the subject showed a generalized fear response to the NA mouse if we reversed the order of presentations of NA and A mice on day 1, we performed a control experiment to determine whether the preference of the subject for the NA compared to A mouse on day 2 was due to the order of presentation on day 1. We thus presented a group of control subjects with two NA mice in successive sessions on Day 1, with the encounters separated by 30 min, similar to the AD paradigm (control 1). When tested in a memory recall test on day 2, the control group mice showed no preference between the two familiar NA stimulus mice, indicating that there was no memory of the order of presentation after 24 hours (Fig. 1b).

It is also possible that the preference of the subjects for the NA compared to A mouse on day 2 may not be driven by the memory of the subject’s prior experiences on day 1 but may rather result from differences in the intrinsic behaviors displayed by the NA versus A mice under the cups on day 2. We therefore performed a second control experiment in which we exposed a group of naïve subjects to a novel NA mouse and a novel prescreened A mouse confined to wire cup cages as in the AD recall test, but without prior encounters with the subject (control 2). These control subjects also showed no significant preference for the NA compared to A mouse (Fig. 1b). Furthermore, the SDI was significantly greater in the test group of subjects than in both control groups (one-way ANOVA with post hoc Tukey test: F(2, 42) = 9.512, p = 0.0004; test vs. control 1, p = 0.01; test vs. control 2, p = 0.0004; Fig. 1c). The test group also exhibited a smaller total interaction time with the NA and A mouse than did the control 1 group (p = 0.003; Fig. 1d), which was driven by the social avoidance towards the A mouse in the test group.

These results suggest that the subjects remembered the different valences of the social interactions with the NA compared to A mouse for at least 24 h. Social memory is believed to have at least two distinct components, the assessment of familiarity versus novelty (social novelty recognition memory, SNRM) and the recollection of the prior experiences with individual conspecifics. Our studies of AD indicate that our subjects could recollect the valence of their social encounters over a 24 h period. To assess whether the mice may also form a lasting memory of familiarity, we performed an experiment in which a subject encountered an NA mouse on day 1 in a single 10 min session. In a SNRM recall session on day 2, we presented the subject with the previously encountered, now-familiar NA mouse and a second novel NA mouse confined to the wire cup cages. The subjects spent significantly more time investigating the novel NA mouse compared to the familiar NA mouse, supporting retention of SNRM at 24 (Supplementary Fig. 2, ctrl-3). Thus, under the conditions of our experiment, mice show both a long-term familiarity memory and long-term social valence recollection memory.

### Avpr1b-expressing neurons in dCA2 are necessary for the formation of AD-STDM

To determine whether dCA2 neurons are required for AD-STDM after our AD paradigm, we used a chemogenetic approach to silence dCA2 pyramidal neurons by injecting AAV-DIO-hM4Di-mCherry into dCA2 of *Avpr1b-Cre*^+/−^ (Cre) mice, an approach that successfully targets nearly all dCA2 pyramidal neurons. The same virus was injected in CA2 of *Avpr1b-Cre*^−/−^ (WT) littermate controls. We first silenced dCA2 Avpr1b-expressing neurons on Day 1 by injecting mice with the hM4Di ligand Clozapine N-oxide (CNO) 30 min prior to the day 1 learning session. We then assessed social discrimination memory during the Day 2 recall session in the absence of CNO, when dCA2 should be active (Fig. 2a–b).

**Figure 2.**
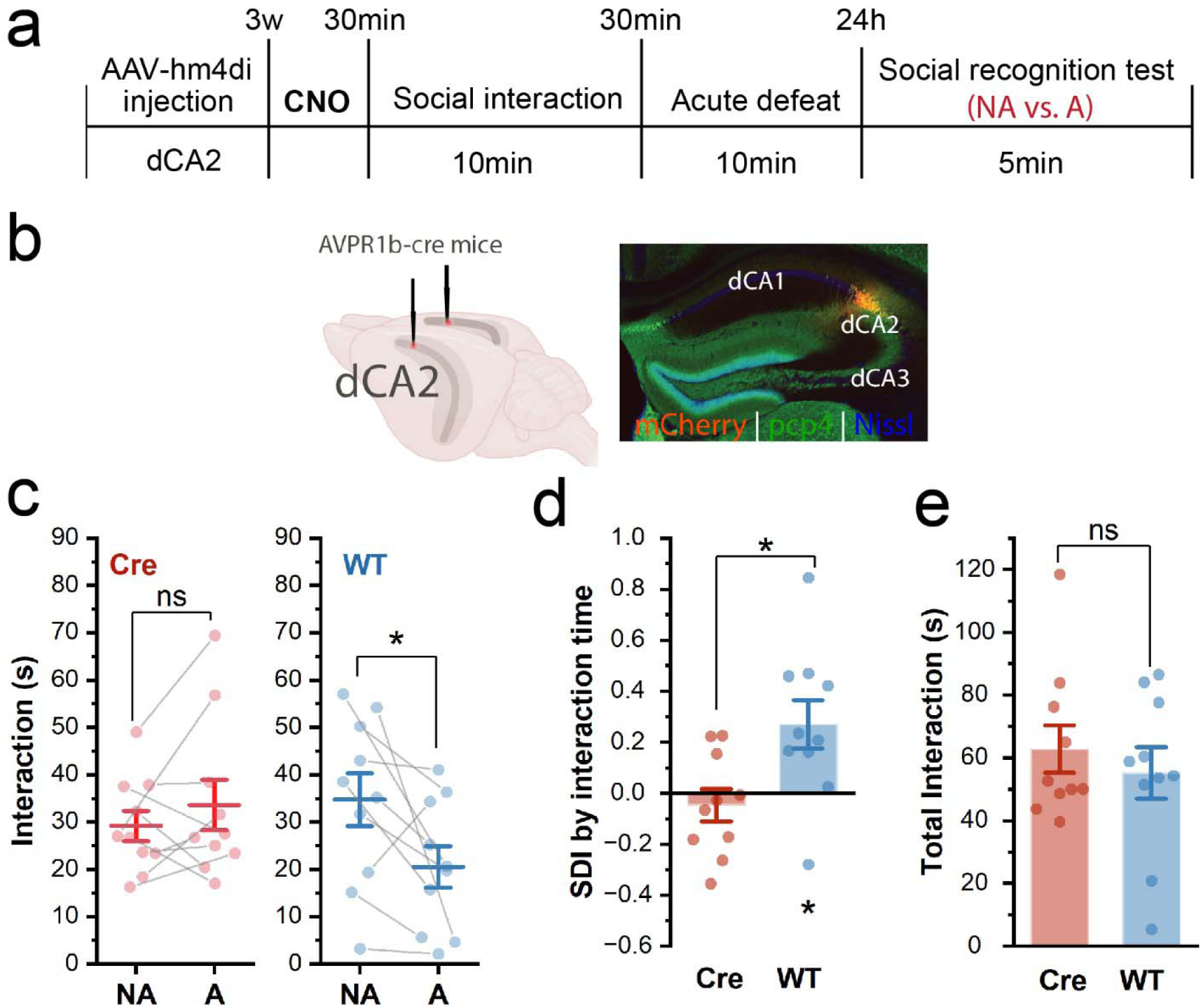
Chemogenetic inhibition of Avpr1b-expressing dCA2 neurons impairs the acquisition of AD-STDM. **a.** Experimental timeline. AAV-DIO-hM4Di-mCherry was injected into dCA2 of *Avpr1b-Cre*^+/−^ mice (n = 10) and wild-type littermate (WT) controls (n = 9). CNO was administered 30 min before the day 1 learning session. Subject mice then went through the AD-STDM paradigm. **b.** Schematic of the viral injection into dCA2 and a representative slice showing hM4Di-mCherry expression in dCA2 of an *Avpr1b-Cre*^+/−^ mouse. **c.** Direct interaction time during the AD-STDM test. Following CNO administration, *Avpr1b-Cre*^+/−^ mice showed no difference in interaction time between the familiar NA and A (blue), whereas WT controls spent more time interacting with the NA than with the A (red). paired t test, Cre: t = −1.028, p = 0.331; WT: t = 2.449, p = 0.037. **d.** Social discrimination index calculated from the direct interaction time were lower in *Avpr1b-Cre*^+/−^ mice than in WT controls. two-sample t test, t = −3.86, p = 0.001; one-sample t test vs. 0: Cre t = −0.75, p = 0.47; WT t = 2.84, p = 0.02. **e.** Total interaction time did not differ between the Cre and WT groups. Two-sample t test, t = 0.674, p = 0.51 ns, not significant, *p < 0.05

Silencing dCA2 Avpr1b-expressing neurons did not change the behavioral properties of the subjects during interactions with the NA or A mice on day 1 (Supplementary Fig. 3). Similar to the WT group, the Cre group showed a longer duration of immobility, a higher maximum movement speed, and a longer occupancy of edge zones of the arena during the AD session with the A mouse compared to the interaction session with the NA mouse (Supplementary Fig. 3a–c). Furthermore, on day 1, the number of attacks received and active investigation times with the two stimulus mice did not differ between the Cre and WT groups (Supplementary Fig. 3d–h).

However, silencing dCA2 in *Avpr1b-Cre*^+/−^ mice on Day 1 disrupted the preference for the NA mouse during the Day 2 recall session. Thus, the WT mice spent significantly more time interacting with the familiar NA than with the A, whereas *Avpr1b-Cre*^+/−^ mice showed no preference. Similarly, SDI was significantly lower in *Avpr1b-Cre*^+/−^ mice than in WT controls (Fig. 2d), with total interaction time unchanged (Fig. 2e). Together, these results support the necessity of Avpr1b-expressing dCA2 neurons during AD-STDM formation. This finding is consistent with previous work that dCA2 plays an essential role in discriminating a social fear-conditioned mouse from a non-threatening mouse^22^.

### Avpr1b-expressing neurons in dCA2 are required for the recall of AD-STDM

Given previous studies of the importance of dCA2 on the recall phase of SNRM^8,22^, we next examined whether dCA2 is also required during the recall of AD-STDM. To this end we administered CNO 30 min before the recall test in the Cre and WT mice that had received dCA2 injections of AAV-DIO-hM4Di-mCherry (Fig. 3a). WT mice preferentially investigated the familiar NA, whereas Cre mice showed no significant interaction preference. SDI was significantly lower in Cre mice than in WT controls. SDI was also significantly above zero in WT mice but not in *Avpr1b-Cre*^+/−^ mice. Total interaction time did not differ significantly between groups (Fig. 3d).

**Figure 3.**
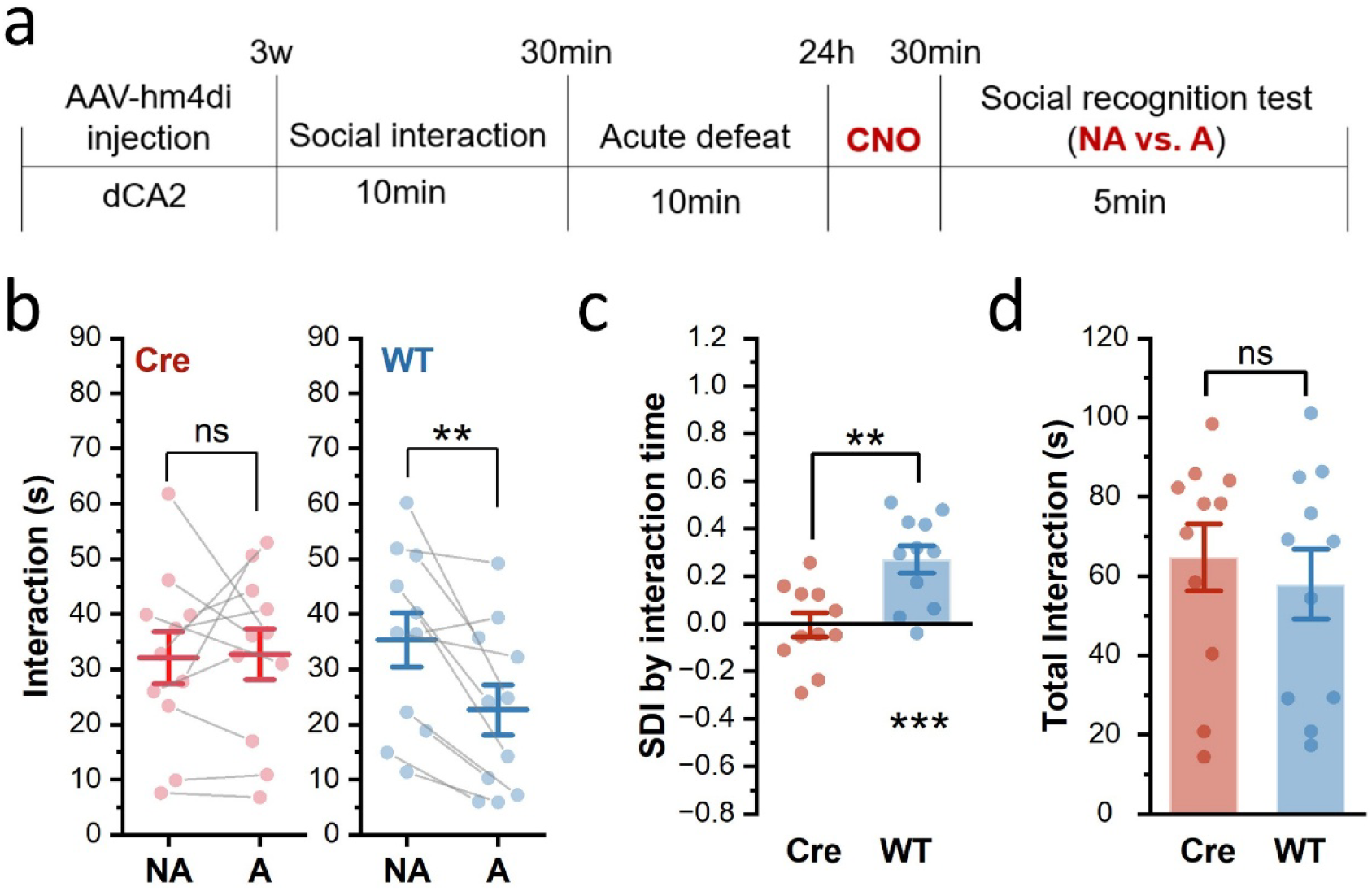
Chemogenetic inhibition of Avpr1b-expressing dCA2 neurons impairs the recall of AD-STDM. **a.** Experimental timeline. AAV-DIO-hM4Di-mCherry was injected into dCA2 of *Avpr1b-Cre*^+/−^ mice and Cre-negative wild-type (WT) littermate controls. Three weeks after viral injection, subject mice underwent the AD-STDM paradigm. CNO was administered 30 min before the 5-min social-recognition test. **b.** Direct interaction time during the social-recognition test. Following CNO administration, *Avpr1b-Cre*^+/−^ mice showed no difference in interaction time between the familiar NA and A (left), whereas WT controls spent more time interacting with the NA than with the A (right). (n=11 in each group; paired t test, Cre: t = −0.16, p = 0.88; WT: t = 3.73, p = 0.003 **c.** SDI was lower in *Avpr1b-Cre*^+/−^ mice than in WT controls. (Unpaired t test: t = −3.62, p = 0.002; one-sample t test vs. 0: Cre: t = −0.12, p = 0.91; WT: t = 4.73, p = 0.0008) **d.** Total interaction time did not differ between the Cre and WT groups. Two-sample t test, t = 0.557, p = 0.58 *p < 0.05, **p < 0.01, ***p < 0.001

Thus, inhibiting Avpr1b-expressing dCA2 neurons during recall impaired discrimination between the familiar NA and A without a detectable reduction in total interaction time, supporting their involvement in AD-STDM recall.

### Avpr1b-expressing neurons in vCA2 are necessary for the formation of AD-STDM

The development of AD-STDM depends on three basic factors: 1) recognizing the stimulus mice are familiar; 2) remembering the two rounds of social interactions with different valences that occurred on Day 1; 3) correctly assigning the social experience to the social identity. While Avpr1b-expressing vCA2 neurons are not important for SNRM^12^, whether these neurons are important for memorizing past experiences and associating them with the correct conspecifics is not known. To this end, we first investigated whether Avpr1b-expressing vCA2 neurons contribute to AD-STDM formation. Similar to the dCA2-silencing approach, we injected AAV-DIO-hM4Di-mCherry in vCA2 of *Avpr1b-Cre^+/−^*mice and WT littermates and then injected both groups of mice with CNO 30 min prior to the AD protocol on day 1 (Fig. 4a–b). Similar to the chemogenetic silencing experiment in dCA2, *Cre^+/−^* and WT groups displayed similar responses to the NA and A mice during the Day 1 learning session (Supplementary Fig. 5), suggesting that silencing Avpr1b-expressing vCA2 neurons does not change how animals behave during non-aversive and aversive social interactions.

**Figure 4.**
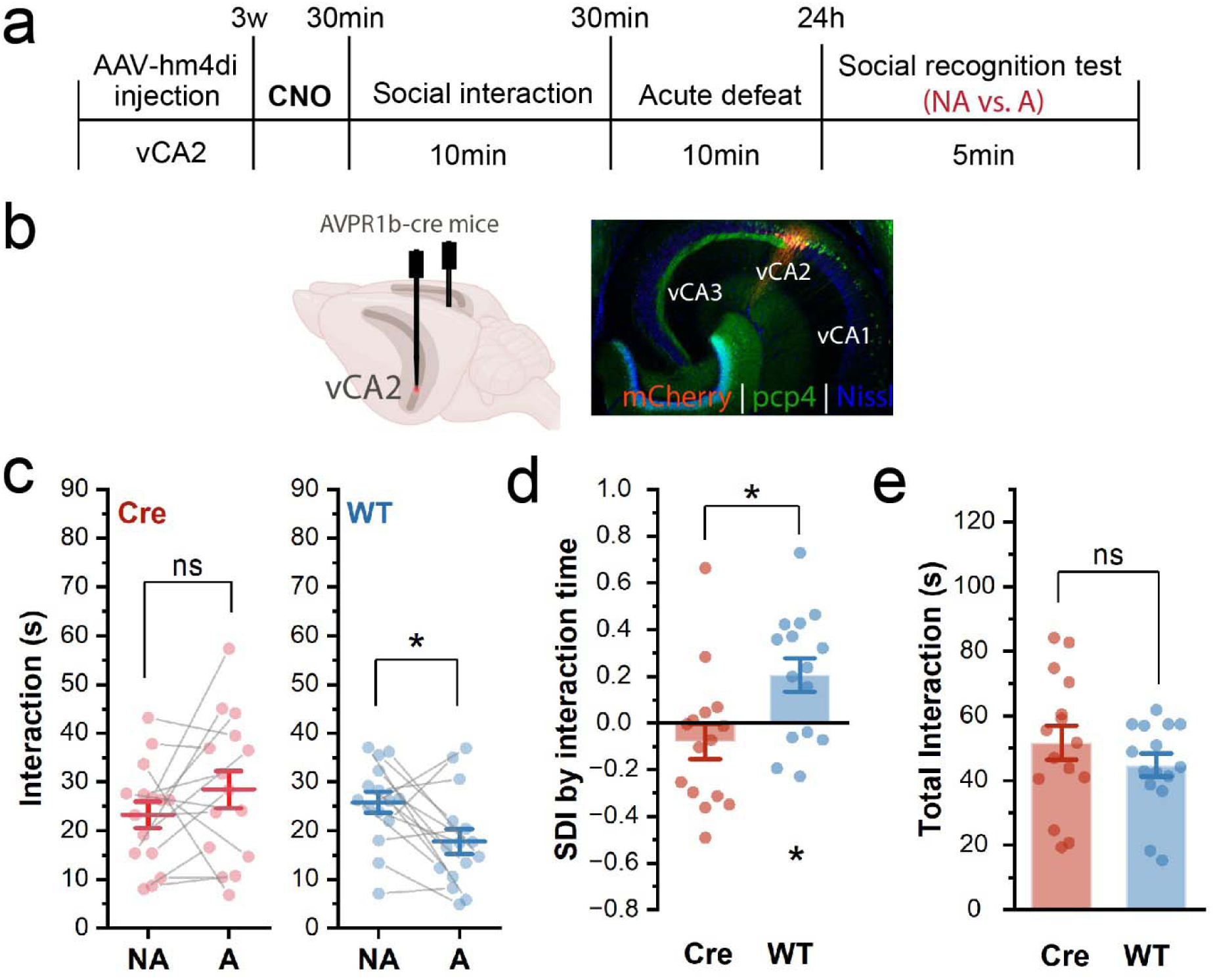
Chemogenetic inhibition of Avpr1b-expressing vCA2 neurons impairs the acquisition of AD-STDM. **a.** Experimental timeline. AAV-DIO-hM4Di-mCherry was injected into vCA2 of *Avpr1b-Cre*^+/−^ mice and Cre-negative wild-type (WT) littermate controls. Three weeks after viral injection, CNO was administered 30 min before the day 1 learning session. Subject mice then underwent the AD-STDM paradigm. **b.** Schematic of the viral injection into vCA2 and a representative image showing hM4Di-mCherry expression in vCA2 of an *Avpr1b-Cre*^+/−^ mouse with PCP4 stained in green. **c.** Direct interaction time during the social-recognition test. *Avpr1b-Cre*^+/−^ mice showed no difference in interaction time between the familiar NA and A, whereas WT controls spent more time interacting with the NA than with the A. (n = 15 per each group; paired t test, Cre: t = −1.325, p = 0.21; WT: t = 2.53, p = 0.023) **d.** SDI was lower in *Avpr1b-Cre*^+/−^ mice than in WT controls. (two-sample t test, t = −2.751, p = 0.01; one-sample t test vs. 0, Cre t =-1.05, p = 0.312; WT t = 2.9, p = 0.012.) **e.** Total interaction time did not differ between the Cre and WT groups. Two-sample t test, t = −1.08, p = 0.29 ns, not significant; *p < 0.05, \*\***p** < 0.01, and \*\*\*\***p** < 0.0001.

During the Day 2 recall test (with no CNO present), WT mice displayed normal AD-STDM, spending significantly more time investigating the NA than the A mice. In contrast, the CA2-silenced *Cre^+/−^* mice showed a complete absence of AD-STDM, with no difference in interaction time with the two stimulus mice and a significantly lower SDI than observed for the WT group (Fig. 4c-d). The total interaction time did not differ significantly between groups (Fig. 4e). Thus, we conclude that the activity of Avpr1b-expressing vCA2 neurons is required for the formation of AD-STDM.

The ventral hippocampus, and particularly ventral CA1, has been shown to modulate anxiety-like behaviors and fear conditioning in rodents^18–21^. To determine if the effect of silencing CA2 on AD-STDM may be due to a change in anxiety-like or fear-like behavior, in general, we evaluated the effect of chemogenetic silencing vCA2 Avpr1b-expressing neurons. We found that silencing vCA2 caused no change in anxiety-like behaviors measured by either the open field test or elevated plus maze test. Thus, there was no difference in either distance traveled or percentage of time spent in the center during the open-field test between WT mice and *Cre^+/−^* mice expressing hM4Di, both injected with CNO (Supplementary Fig. 8a). Similarly, there was no difference in these groups in time spent in the open and closed arms during the elevated plus maze test of anxiety (Supplementary Fig. 8b). In a separate cohort, we examined the effect of silencing CA2 during a fear conditioning test. We injected WT female mice and *Cre^+/−^* female mice expressing hM4Di with CNO 30 minutes prior to exposing the mice to a fear conditioning paradigm in which we paired a tone with foot shocks in a given context^25^. After 24 and 48 hours, we performed auditory and contextual fear recall tests, respectively (Supplementary Fig. 9). We did not detect differences in freezing between two group during either the auditory recall test or the contextual recall test. Together, these findings support a role for Avpr1b-expressing vCA2 neurons in AD-STDM formation, with no detectable effects on non-social anxiety-related measures or on the formation of auditory and contextual fear memory.

### The role of vCA2 Avpr1b-expressing neurons in the recall of AD-STDM

Next, we examined whether Avpr1b-expressing vCA2 neurons contribute to AD-STDM recall by injecting CNO 30 min before the recall test on day 2 in *Cre^+/−^* mice and WT controls that had been injected in vCA2 with AAV-DIO-hM4Di-mCherry (Fig. 5a). WT mice spent significantly more time investigating the familiar NA than the A, whereas the Cre mice showed no preference between the NA and A mice (Fig. 5b). However, there was a significant amount variability in the cohorts and the SDIs did not differ significantly between groups (Fig. 5c). Total interaction time also showed no significant between-group difference (Fig. 5d).

**Figure 5.**
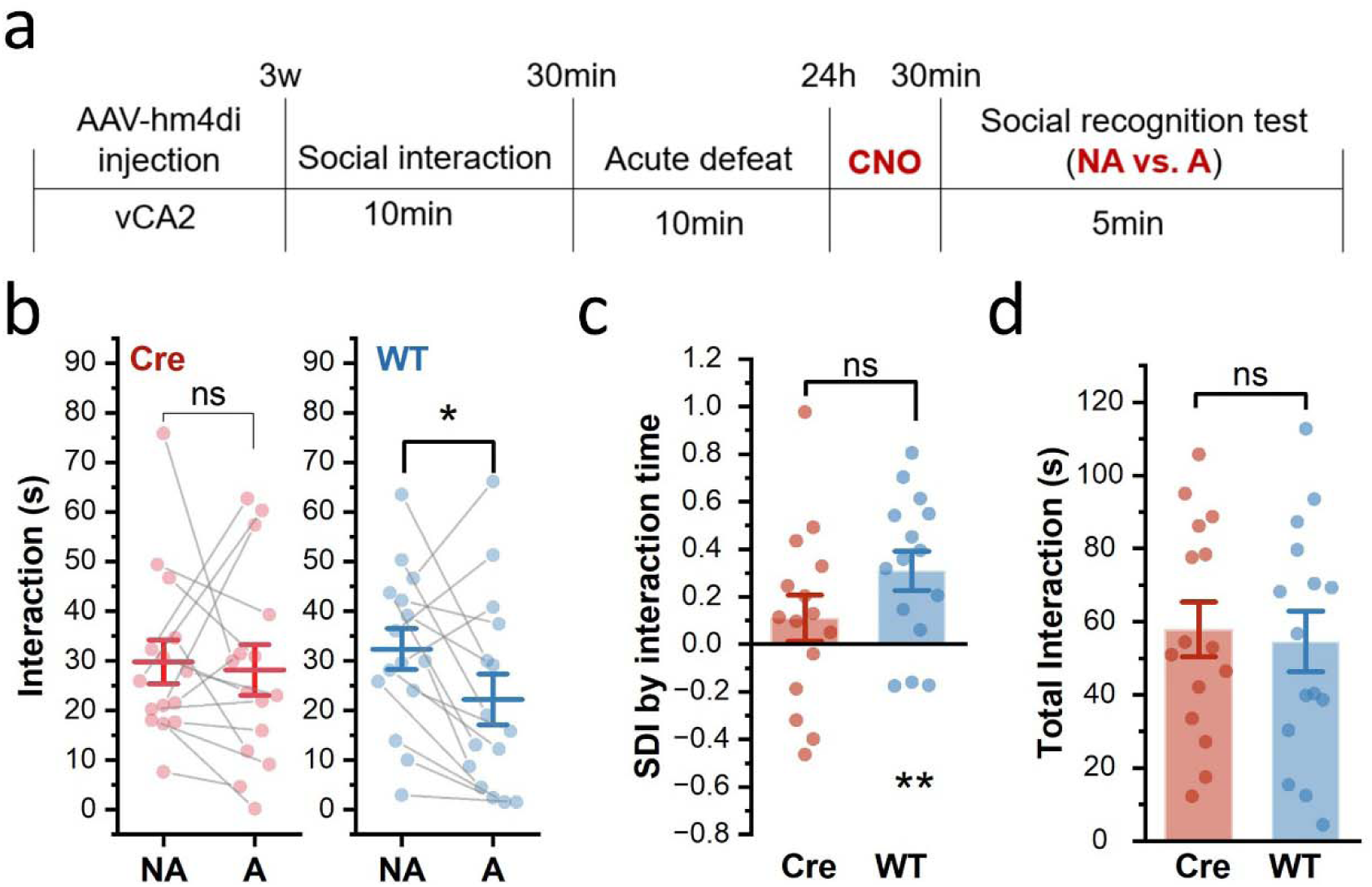
Chemogenetic inhibition of Avpr1b-expressing vCA2 suggests a possible role in recall of AD-STDM. **a.** Experimental timeline. AAV-DIO-hM4Di-mCherry was injected into vCA2 of *Avpr1b-Cre*^+/−^ mice and Cre-negative wild-type (WT) controls. Three weeks after viral injection, subject mice underwent the AD-STDM paradigm. CNO was administered 30 min before the 5-min social-recognition test. **b.** Direct interaction time during the social-recognition test. Following CNO administration, *Avpr1b-Cre*^+/−^ mice showed no significant difference in interaction time between the familiar NA and A, whereas WT controls spent more time interacting with the NA than with the A. (n=15 in each group; paired t test, Cre: t = 0.28, p = 0.78; WT: t = 2.39, p = 0.03) **c.** While only the SDI of the WT group significantly prefers to the NA, SDI did not differ significantly between the Cre and WT groups. (Unpaired t test: t = 1.56, p = 0.129; One-sample t test: Cre: t = 0.28, p = 0.78; WT: t = 2.39, p = 0.03) **d.** Total interaction time did not differ significantly between the Cre and WT groups. Two-sample t test, t = −0.296, p = 0.77 *p < 0.05 and **p < 0.01.

To exclude the possibility that the behavioral differences between *Cre^+/−^* and WT groups in AD-STDM storage or recall are due to genetic background differences (e.g., expression of Cre) rather than the chemogenetic silencing of CA2 neurons, we carried out a separate set of control experiments. We injected AAV-DIO-hM4Di-mCherry in vCA2 and dCA2 in separate cohorts of *Avpr1b-Cre*^+/−^ and WT mice. However, rather than injecting the mice with CNO, we injected cohorts of mice with saline, both 30 min before the AD learning session on Day 1 and 30 min before the recall session on Day 2 (Supplementary Fig. 7a). Similar to the other WT control groups, the saline-treated mice all showed significant AD-STDM, with a clear preferential interaction with the NA compared to A mice during the recall session; there was no difference between genotypes in either SDI or total interaction time (Supplementary Fig. 7).

### Effect of social interactions on cFos expression in dCA2 and vCA2

To determine how CA2 neurons respond to social encounters during AD, we quantified expression of the immediate-early gene cFos as a marker of neuronal activity one hour after either a non-aggressive social interaction (SI) or AD and compared this to cFos levels in singly housed home-cage controls (HC) (Figure 6). In dCA2, we observed a significant increase in the number of cFos+ cells relative to HC levels after both SI and AD (Fig. 6a, d), with no significant difference between SI and AD. In vCA2 we also observed a significant increase in the number of cFos+ cells after both SI and AD. However, in contrast to dCA2, the number of cFos+ cells in vCA2 was significantly greater after AD compared to SI (Fig. 6b, g). We also found a greater number of cFos+ cells in vCA2 compared to dCA2 under all conditions (Supplementary Figure 11a, b).

**Figure 6:**
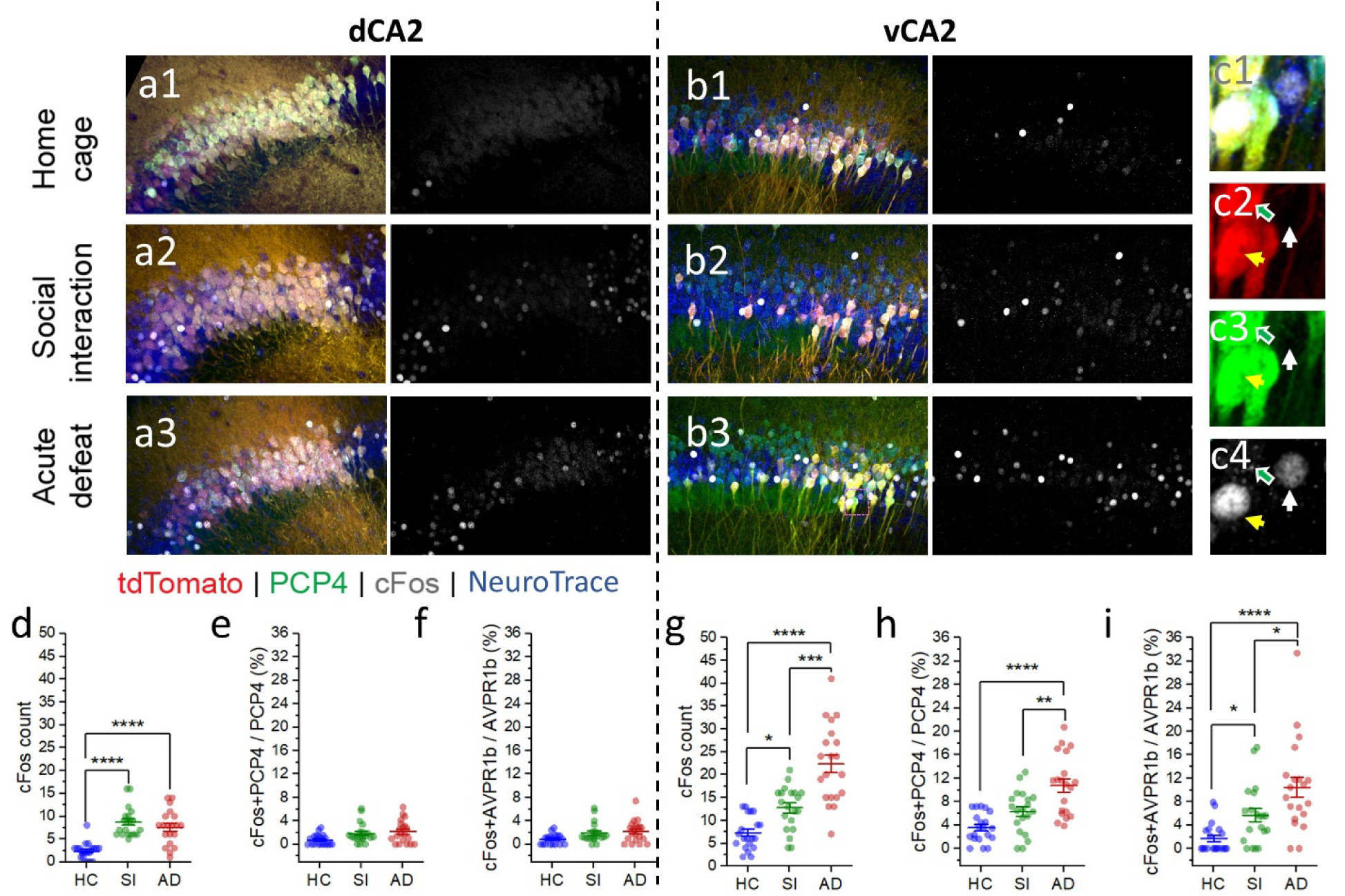
Acute defeat increases cFos expression in vCA2 compared with home-cage or non-aggressive social-interaction conditions. **a–b.** Representative hippocampal slices containing dCA2 (a1–a3) and vCA2 (b1–b3) from Avpr1b-tdTomato mice in the home-cage (HC), 10-min social-interaction (SI), and 10-min acute-defeat (AD) groups. Avpr1b-tdTomato is shown in red, PCP4 in green, cFos in gray, and NeuroTrace in blue. **c.** High-magnification view of the magenta dotted box in b3. The yellow arrow indicates an example cell co-labeled for cFos, PCP4, and Avpr1b. The green arrow points an example cell expressing PCP4 and Avpr1b, but not cFos. The white arrow indicates an example cell expressing cFos, but not PCP4 or Avpr1b. **d–f.** In dCA2, the total number of cFos was higher in the SI and AD groups than in the HC group (**d**). No differences were observed in the percentage of PCP4+ neurons expressing cFos (**e**) or the percentage of Avpr1b-expressing neurons expressing cFos (**f**). (Group effects: **d**, χ²(2) = 15.753, p = 0.000380; **e**, χ²(2) = 1.519, p = 0.4679; **f**, χ²(2) = 1.548, p = 0.4611; Two-sided Wald z-tests with Holm correction: **d**, HC vs. SI, p < 0.0001; HC vs. AD, p < 0.0001; SI vs AD, p = 0.947) **g–i.** In vCA2, the total number of cFos was higher in the SI group than in the HC group and was further increased in the AD group relative to both the HC and SI groups (**g**). The percentages of PCP4+ neurons (**h**) and Avpr1b-expressing neurons (**i**) expressing cFos were also higher in the AD group than in both the HC and SI groups. (Group effects: **g**, χ²(2) = 17.833, p = 0.000134; **h**, χ²(2) = 14.344, p = 0.000768; and **i**, χ²(2) = 17.054, p = 0.000198. Two-sided Wald z-tests with Holm correction: **g**, HC vs. SI, p = 0.0467; HC vs. AD, p < 0.0001; SI vs. AD, p = 0.00097; **h**, HC vs. SI, p = 0.0951; HC vs. AD, p < 0.0001; SI vs. AD, p = 0.009; **i**, HC vs. SI, p = 0.024; HC vs. AD, p < 0.0001; SI vs. AD, p = 0.012.) n = 4 mice and 20 slices per group, with five slices per mouse. Coordinates relative to bregma were −1.34 to −2.06 mm for dCA2 and −3.72 to −4.44 mm for vCA2. Data were analyzed using a mixed-model analysis. *p<0.05, **p<0.01, ***p<0.001, ****p<0.0001

As the CA2 region contains a mixture of excitatory pyramidal neurons and inhibitory neurons, we explored how cFos levels depend on the molecular identity of CA2 cells by co-staining for cFos and the CA2 pyramidal cell marker PCP4 in *Avpr1b-Cre x Ai14* reporter mice. We previously found that PCP4 and Avpr1b are co-expressed in nearly all pyramidal neurons in the dorsal CA2 region of the hippocampus. In ventral CA2, approximately 60% of neurons express PCP4 whereas 40% of neurons express Avpr1b, all of which co-express PCP4^12^. Although the dCA2 pyramidal layer contained more PCP4-positive neurons than vCA2, both in absolute counts (Supplementary Fig. 10) and as a percentage of NeuroTrace-positive cells^12^, we found very few cFos-expressing cells in PCP4-positive or tdTomato-positive dCA2 pyramidal neurons, with no significant difference between HC cFos-expressing cell counts and those after SI or AD (Fig. 6e-f). In contrast, vCA2 showed a higher number of cFos-expressing cells that expressed PCP4 or tdTomato compared to dCA2 under all conditions (Supplementary Figure 11c-f).

Moreover, there was a significant increase in the number of PCP4-positive and tdTomato-positive neurons that expressed cFos following SI and AD compared to HC levels, with cFos-expressing cell counts significantly greater after AD compared to SI (Fig. 6h-i).

## Discussion

Our findings demonstrate a critical role of dorsal and ventral hippocampal CA2 in social threat discrimination memory of an aversive social encounter during a single episode of acute social defeat with one conspecific compared to a non-aversive social encounter with a different conspecific. Under the acute defeat-associated social threat discrimination memory (AD-STDM) paradigm, mice preferentially investigated the non-aggressor mouse over the aggressor mouse 24 h after the encounters with both individuals.

Chemogenetic inhibition of Avpr1b-expressing pyramidal neurons in either dCA2 or vCA2 during the AD-STDM learning trials led to a significant impairment in AD-STDM assessed during the 24 h discrimination recall trial. We also found that selective chemogenetic inhibition of dCA2 during the AD-STDM recall trial also significantly impaired AD-STDM. However, the evidence for the importance of vCA2 in AD-STDM recall is less clear-cut. Thus, the mice in the group in which vCA2 was silenced during the recall trial failed to show a normal preference for interacting with the non-aggressor mouse compared to the aggressor mouse. Although, the mean preference index of the vCA2-silenced group was less than that of the control group, the difference was not statistically. This may reflect the large variance in preference index in the two groups of mice in this set of experiments. Thus, whereas our data clearly support a role for dCA2 and vCA2 in the acquisition of AD-STDM, our results on AD-STDM recall provide strong support for a requirement for dCA2 and less clear support for a role of vCA2.

A role of dCA2 and vCA2 in AD-STDM from our chemogenetic studies is supported by experiments using cFos staining as a measure of neural activation during the AD-STDM paradigm. Thus, we found that cFos expression is increased in neurons in both dCA2 and vCA2 during both non-aggressive and aggressive social interactions to levels significantly above those observed in the home cage. Of further interest, whereas cFos levels in dCA2 were increased to similar levels by the aggressive and non-aggressive social interactions, cFos levels in vCA2 were significantly greater following acute defeat compared to non-aggressive social interaction. Thus, we conclude that not only are dCA2 and vCA2 required for AD-STDM, but that vCA2 may also signal the valence of social interaction. Together, these findings establish a contribution of CA2 along the dorsoventral axis to AD-STDM formation and extend our understanding of how CA2 supports social memory following aversive encounters.

The AD-STDM paradigm complements existing approaches to studying memory for socially relevant threats. Previous work from our laboratory initially demonstrated that dCA2 is required for discrimination between conspecifics associated with or without foot shocks in a social fear-conditioning paradigm^22^. Later work from Zorab et al.^23^ confirmed a role of dCA2 in social threat memory, and expanded this to include discrimination of an aggressor following acute defeat compared to a non-aggressor following a non-threatening encounter. However, this study used a memory interval of only 5 min, and so its relevance to hippocampal-dependent long-term memory storage is uncertain. Moreover, the CD-1 male and CFW female stimulus strains are larger-bodied than the C57BL/6 subject mice, raising the possibility that size-related cues contributed to generalized avoidance. In our study, we assessed discrimination memory of an aggressor versus a non-aggressor mouse using male BALB/c stimulus mice that were age- and size-comparable to the C57BL/6 subject mice. Furthermore, we used a 24 hour memory interval, consistent with typical time frames known to require the hippocampus.

Our experiments also complement studies identifying a role for vCA1 in social valence^23,24^. Zorab and colleagues demonstrated contributions of serotonergic and neurotensinergic inputs to vCA1 in the encoding of the memory of positive and negative social valence, respectively, 5-min following the initial interactions^23^. Suto and colleagues identified a critical role of synaptic plasticity in a vCA1 to basolateral amygdala circuit in updating the valence of familiar individuals^24^. Our study extends these prior results to provide the first comparison of the contributions of dorsal and ventral CA2 to the formation of long-term social threat discrimination memory after a single episode of social defeat.

The contribution of vCA2 to social threat discrimination is of particular interest in light of our previous finding that inhibition of vCA2 did not impair SNRM^12^. This suggests that vCA2 is of selective importance in social recollection memory compared to social familiarity memory. In contrast, participation of dCA2 is required for both social familiarity and recollection of the valence of a prior social encounter. Also in contrast to vCA1, we found that inhibition of vCA2 did not alter anxiety-related behaviors or auditory and contextual non-social fear memory. Thus, similar to dCA2, vCA2 appears to be particularly important for social forms of memory. However, as vCA2 comprises a heterogeneous population of neurons, only 40% of which express Avpr1b, it is possible that an Avpr1b-negative population of vCA2 neurons contributes to non-social behaviors.

In principle, Avpr1b-expressing vCA2 neurons may contribute to social threat discrimination memory by encoding the association of social experiences of different valences with the particular identity of the conspecifics. Alternatively, vCA2 may selectively encode the valence of a social interaction and the association with social identity may be carried out by a downstream region, such as vCA1, that receives social identity input from dCA2 and social valence input from vCA2. A third possibility is that vCA2 neurons may simply participate in recollection memory of the social identities of the conspecifics, with information about valence contributed from other brain regions. Although further studies will be required to distinguish these possibilities, our cFos findings that vCA2 is activated more strongly by an episode of acute defeat than by an episode of non-aggressive social interaction favors the idea that vCA2 participates in the encoding of social valence, not simply identity.

Candidate circuits underlying the contributions of dCA2 and vCA2 to AD-STDM are suggested by consideration of the brain regions from where these areas receive input and provide output. Our previous work showed that dCA2 preferentially targets vCA1a, the area closest to ventral subiculum, and dorsal anterior lateral septum (LS), whereas vCA2 preferentially targets vCA1b/c and ventral posterior LS^12^. In view of the involvement of vCA1 in social valence^23,24^ and LS in social avoidance following social defeat^26^, these complementary projections raise the possibility that dorsal and ventral CA2 influence different components of downstream processing during aversive social learning. Furthermore, upstream neuropeptide signaling may also provide a level of regulation. Avpr1b expression extends along the CA2 dorsoventral axis, and Oxtr expression has been reported in dorsal and ventral CA2/distal CA3^9,12,27^. Projection-specific manipulations during learning and recall will be needed to determine how these anatomically distinct pathways contribute.

One important question about our cFos results is the identity of the cells that are activated by the aggressive and non-aggressive social interactions. CA2 comprises roughly 80% pyramidal neurons and 20% inhibitory neurons. Moreover, whereas pyramidal neurons in dCA2 comprise a more-or-less uniform population, with nearly all neurons co-expressing all standard CA2 markers, including RGS-14, STEP, PCP4 and Avpr1b-expression, neurons in vCA2 express a subset of markers expressed in dCA2^12^. Thus, vCA2 shows only weak expression of RGS-14 and STEP. Furthermore, the dCA2 markers that are expressed in vCA2 are more heterogeneous, with 60% of vCA2 neurons expressing PCP4 and 40% expressing Avpr1b (all of which express PCP4), the latter based on tdTomato expression in an *Avpr1b-Cre x Ai14* reporter mouse.

We thus examined cFos staining in subsets of pyramidal neurons by co-staining for PCP4 or tdTomato (as a marker of Avpr1b-expressing cells). Although, as noted above, cFos levels were elevated about homecage levels in dCA2 as a whole, when we restricted our analysis to dCA2 pyramidal neurons, we found no effect on cFos expression following SI or AD, using either PCP4 or tdTomato markers to identify dCA2 pyramidal cells. In contrast, we found significant increases in cFos levels in the PCP4 and Avpr1b-expressing pyramidal cell populations in vCA2. Moreover, in both populations, we observed a significantly greater increase in cFos levels following AD compared to the non-aggressive social interaction. Future experiments will be needed to determine whether the cFos+ cells that fail to express CA2 pyramidal cell markers may represent a population of inhibitory neurons.

The lack of cFos expression in dCA2 pyramidal neurons is consistent with findings that these cells are normally resistant to the induction of long-term potentiation^28^, which is thought to result, at least in part, from expression of RGS14 to limit postsynaptic calcium signaling, whereas STEP contributes to metabotropic glutamate receptor-dependent long-term depression^29–31^. The absence of cFos induction in dCA2 may therefore reflect limited engagement of plasticity-related mechanisms that, in other cell types, support learning and memory^32^. Our finding of activity-dependent cFos expression in vCA2 pyramidal neurons suggests that these cells may be less resistant to the induction of synaptic plasticity compared to dCA2 neurons.

Together, these findings identify Avpr1b-expressing neurons in dorsal and ventral CA2 as contributors to the formation of social threat discrimination memory following a single AD episode, with an additional demonstrated contribution of dCA2 during recall. The results extend the behavioral role of CA2 beyond that revealed by SNRM and social aggression, and motivate investigation of how CA2 molecular diversity, neuromodulation, and anatomically distinct outputs across the dorsoventral axis support distinct forms of memory for socially significant experiences.

## Methods

### Animals

All procedures were approved by the Columbia University Institutional Animal Care and Use Committee under protocol AABJ7554. All experimental and stimulus mice were male and 2-4 months old except for the fear conditioning experiments where female mice were used. Unless otherwise specified, mice were housed with littermates in groups of two to five per cage. Mice were maintained on a 12-h light/dark cycle under air-filtered, temperature- and humidity-controlled conditions, with food and water available ad libitum.

C57BL/6J mice (The Jackson Laboratory, 000664) were used to develop the AD-STDM paradigm. *Avpr1b-Cre*^+/−^ mice^9^ (The Jackson Laboratory, 036876) and their *Avpr1b-Cre*^−/−^ littermates were used in the chemogenetic silencing experiments. *Avpr1b-Ai14* mice were bred in-house by crossing female *Avpr1b-Cre*^+/−^ mice with male Ai14 mice (The Jackson Laboratory, 007914) and were used for cFos quantification. Age-matched male BALB/cJ mice (The Jackson Laboratory, 000651) were used as aggressors (A), while age-matched male BALB/cByJ mice (The Jackson Laboratory, 001026) were used as non-aggressor (NA) stimulus mice.

Unless otherwise specified, behavioral experiments were conducted during the light phase under low-intensity red illumination. All behavioral sessions were recorded using a FireWire camera (DMK 31AF03-Z2; The Imaging Source) triggered through ANY-maze software.

### AD-STDM paradigm

Acute defeat-associated social threat discrimination memory (AD-STDM) paradigm was developed to assess long-term memory associating social identity to the social valence of a prior social encounter. One day before the learning phase, subject mice were habituated for at least 10 min to an empty three-chamber apparatus (each chamber: 19 × 40.5 × 22 cm). During the learning phase (Day 1), subject mice were habituated to the testing room for 30 min and then allowed to interact freely with a NA BALB/c in a clean cage for 10 min.

Following a 30-min inter-session interval, each subject mouse was introduced into the home cage of a prescreened A BALB/c mouse for 10 min for a resident-intruder test^33^. The interaction was terminated if continuous attack persisted for 2 min. Test mice were excluded from the analysis if 1) NA BALB/c mice attacked test mice, 2) test mice attacked NA BALB/c mice, or 3) A BALB/c mice failed to attack test mice. The defeat (initiated with a bite and ended when the aggressor stopped biting or chasing the subject) and social investigation (nose-to-nose and nose-anogenital sniffing) were manually annotated. Seven behavioral aspects were analyzed with ANY-maze (Stoelting Co.) during the Day 1 sessions: immobility time, maximal speed, time spend in the edge zone (defined as the space that are within 4 cm from the walls of a 16 x 36 cm cage), attack onset, attack duration, attack bouts, and interaction time. 24 hours later (Day 2), subject mice were habituated to the testing room for 30 min and then placed for 10 min in the three-chamber apparatus, with one empty wire cup located in each side chamber before the recall session. For the AD-STDM test, the familiar NA and A stimulus mice were presented simultaneously, with each mouse confined under a wire cup. The locations of the NA and A mice were randomized and counterbalanced across subjects. Time spent investigating each stimulus mouse (manually annotated direct cup sniffing) 5-min test was quantified and social discrimination index (SDI) was calculated as SDI = (t_NA_ - t_A_) / (t_NA_ + t_A_). To control for a potential effect of encounter order (control-1), separate subject mice interacted sequentially with two NA mice on Day 1 and were tested with the same two mice on Day 2.

Since the first-presented NA was postulated to be ‘more novel’, SDI was calculated as SDI = (t_NA1_ – t_NA2_) / (t_NA1_ + t_NA2_). To determine whether the innate behavioral differences between NA and A mice during the recall test account for the subject’s preference (control-2), naïve subject mice were presented with a novel pair of NA and A mice during the recall test. To confirm that social recognition memory persisted for 24 h (control-3), separate subject mice interacted with one NA mouse on Day 1 and were presented with the familiar NA mouse and a novel NA mouse on Day 2. In these cases, SDI = (t_novel_ – t_familiar_) / (t_novel_ + t_familiar_). The habituation, test duration, and stimulus placement were all identical to the main test protocol. A stimulus mouse could be re-used with a different test mouse only if the new learning session occurred at least 3 days after its previous recall session.

To prescreen aggressors, BALB/cJ mice were singly housed for at least 7 d. During a single screening session, a male C57BL/6J stimulus mouse, 2-4 months old, was introduced into the home cage of an aggressor candidate for a 5-min free-interaction session. A candidate was classified as an A stimulus mouse if it initiated an attack within 3 min and attacked the intruder more than once during the 5-min session.

### Viral injection

*Avpr1b-Cre*^+/−^ and *Avpr1b-Cre*^−/−^ mice were anesthetized with isoflurane (5% for induction and 2-3% for maintenance). Immediately after induction of anesthesia, mice received carprofen (5 mg/kg, i.p.) for analgesia. Mice were then secured in a stereotaxic frame, and a craniotomy was made above each target site. A glass pipette containing the Cre-dependent hM4Di virus AAV2/8-hSyn-DIO-hM4D(Gi)-mCherry (1.9 × 10^12^ pp/mL) was lowered to the target depth. A Nano-Inject II system (Drummond Scientific) was used to deliver 230 nL of virus per hemisphere as 23-nL aliquots at 10-s intervals. Following the completion of each injection, the pipette was left in place for 7 min, and was left at 0.2 mm dorsal to the target for another 5 min before being fully retracted. The following stereotaxic coordinates were used relative to bregma (in mm): dCA2, AP −2.0, ML ±1.8, and DV −1.6; and vCA2, AP −3.0, ML ±3.2, and DV −3.8.

### Dorsal and ventral CA2 chemogenetic silencing

Behavioral experiments were conducted 3 weeks after viral injection. Before testing, subject mice were handled for 6 consecutive days and habituated to intraperitoneal sterile saline injections for 4 consecutive days. Subjects were weighed 1 day before testing to calculate the appropriate Clozapine N-oxide (CNO) dose. CNO (Cayman Chemical item 16882) was prepared in sterile saline on the day of the experiment at a concentration of 1 mg/ml. CNO was administered intraperitoneally at 5 mg/kg 30 min before the relevant behavioral session. In the main chemogenetic AD-STDM experiments, both *Avpr1b-Cre^+/−^*and *Avpr1b-Cre^−/−^* mice received CNO on the assigned learning or recall day (Fig. 2-5). Genetic control mice received saline at the same volume (5 mL/kg) on both Day 1 and Day 2 (Supplementary Fig. 7).

### Immunohistochemistry and imaging

Subject mice were transcardially perfused with ice-cold PBS followed by 4% paraformaldehyde (PFA) in ice-cold PBS. Brains were extracted and fixed overnight in 4% PFA at 4°C. Dorsal hippocampal slices were sectioned coronally, whereas ventral hippocampal slices were sectioned horizontally. Sections were cut at a thickness of 60 µm using a Leica VT1000S vibratome. Unless otherwise specified, free-floating sections were permeabilized and blocked for 2 h at room temperature in PBS containing 5% goat serum and 0.4% Triton X-100. Sections were then incubated overnight at 4°C with primary antibodies diluted in PBS containing 5% goat serum and 0.1% Triton X-100. On the second day, sections were washed three times for 10 min each in PBS and incubated with secondary antibodies for 3 h at room temperature. Sections were then incubated with DAPI (1:1,000; Thermo Fisher Scientific, D1306) in PBS for 10 min at room temperature, followed by three washes in PBS. Sections were mounted using Fluoromount (Sigma-Aldrich) and imaged with a Zeiss LSM 700 confocal microscope.

To validate hM4Di-mCherry expression and its localization within CA2 in *Avpr1b-Cre*^+/−^ mice, sections were labeled with rabbit anti-PCP4 (1:300; Sigma-Aldrich, HPA005792, RRID: AB_1855086) and mouse anti-RFP (1:500; Thermo Fisher Scientific, MA5-15257). The corresponding secondary antibodies were goat anti-rabbit IgG and goat anti-mouse IgG (1:500; Life Technologies). Mice were excluded from the analysis if <50% Avpr1b neurons express mCherry. Similar to the previous method^12^, we estimate <36.8% dCA2 or <29.9% vCA2 pyramidal neurons as the cut-off threshold of exclusion.

For cFos labeling, sections were permeabilized and blocked for 2 h at room temperature in PBS containing 10% goat serum and 1% Triton X-100. Primary and secondary antibodies were diluted in PBS containing 5% goat serum and 0.3% Triton X-100. Sections were incubated overnight at 4°C with guinea pig anti-cFos (1:5,000; Zuckerman Institute Antibody Core) and rabbit anti-PCP4 (1:300; Sigma-Aldrich, HPA005792, RRID: AB_1855086). On the next day, sections were brought to room temperature for 1 h and washed three times in PBS. Sections were then incubated for 3 h at room temperature with goat anti-guinea pig IgG and goat anti-rabbit IgG secondary antibodies (1:500; Life Technologies). For the AD cFos experiments, NeuroTrace 435/455 (1:200, Thermo Fisher Scientific, #N21479, RRID: AB_2572212) was included during secondary-antibody incubation. Following secondary-antibody incubation, sections were incubated with DAPI (1:1,000; Thermo Fisher Scientific, D1306) in PBS for 10 min at room temperature.

### AD cFos quantification

*Avpr1b-Ai14* mice were randomly separated to groups experienced 10-min AD, 10-min non-aggressive social interaction, or the home cage group where mice were singly housed. Each brain was separated into hemispheres. One hour after each behavior, or after 1 h of single housing for home-cage controls, mice were perfused for cFos staining. 60-µm ventral hippocampal slices were obtained by sectioning one hemisphere horizontally (n = 4 mice/group, 7 slices/mouse for Supplementary Fig. 11, 5 slices/mouse for the rest cFos figures), whereas 60-µm dorsal hippocampal slices were obtained by sectioning the counterpart coronally (n = 4 mice/group, 5 slices/mouse). CA2 images were obtained as 5 consecutive z-stack images with a delta of 2.04 µm were selected. After obtaining the images, slices at the following coordinates (relative to the bregma in mm) were selected for quantification DV for dCA2: −1.34, −1.52, −1.7, −1.88, −2.06; DV for vCA2: −3.72, −3.9, −4.08, −4.26, −4.44, and additional −3.36 and −3.54 for supplementary Fig. 11. Images were then assigned a random unique number across all animals, so that the cFos-expressing neurons could be manually quantified blindly to the group or z-plane. The CA2 region of interest was defined as a trapezoid with parallel boundaries of 205 and 325 µm and a perpendicular height of 110 µm as previously reported^12^. Five consecutive optical planes yield a volume of 237,864 µm³ per section. A cFos-expressing neuron was identified as the colocalization with DAPI or NeuroTrace within the same optical plane. Five optical planes were examined together to identify unique cells. FIJI imageJ was used for the quantification.

### Open field test

Following CNO injection, subjects were moved to the testing room and allowed to habituate for 30 min. Mice were then placed in a square arena (50 × 50 cm) for 5 min under bright white light. Following testing, the arena was divided into zones in ANY-maze software: an inner zone (15 × 15 cm) centered in the middle of the arena and an outer zone consisting of the remaining area of the chamber. Time spent and distance traveled in each zone, as well as total distance traveled across both zones, were automatically quantified using ANY-maze tracking software.

### Elevated plus maze

Subject mice were moved to the testing room following CNO injection. After 30 min, subjects were placed in the center of an elevated arena with two enclosed arms, each surrounded by three walls, and two open arms connected by a single plus-shaped platform. Mice were allowed to freely explore the closed and open arms for 5 min under bright white light. ANY-maze software was used to automatically quantify the time spent in the open and closed arms.

### Auditory and contextual fear conditioning test

A 3-day delay fear-conditioning protocol was used to test contextual and auditory fear memory using Med Associates NIR Video Fear Conditioning Chambers. One day before testing, female *Avpr1b-Cre^+/−^*mice and *Avpr1b-Cre^−/−^* WT mice were habituated to the testing room for 1 h. On the training day, mice were injected with CNO (5 mg/kg) 30 min before testing and moved to the testing room. The test box was a sound-attenuated enclosure (59.69 × 71.12 × 31.75 cm) configured with transparent walls, a steel grid floor, lighting directly above the chamber, and no odor (context A). After 150 s, a 5-kHz, 75-dB tone was played for 30 s and co-terminated with a 2-s shock (0.070 mV alternating current). Thirty seconds after the shock, the tone–shock pairing was repeated. This process was repeated once more for a total of three tone–shock pairings. Subjects were then returned to their home cages. Between animals, the chamber was cleaned with 70% ethanol. Twenty-four hours after training, subjects were introduced to a novel context consisting of opaque white walls, opaque white flooring, low-light conditions, and a 1% peppermint odor (context B). Subjects were exposed to five 30-s tones, with a 90-s interval following each tone. Forty-eight hours after training, mice were returned to the original context in which they had received the shocks (context A) for 6 min. Across all days, videos were recorded, and freezing was automatically scored using Med Associates Video Freeze software.

### Statistics

For the behavioral comparisons, data passing the Kolmogorov–Smirnov normality test was analyzed using t-tests or ANOVA with Tukey’s or Šídák’s post hoc test; non-normal data was analyzed using Mann–Whitney or Kruskal–Wallis tests with Dunn’s post hoc test (α = 0.05). Linear mixed-effects models were used for the AD cFos quantification, with experimental group as a fixed effect and mouse identity as a random intercept to account for sections sampled from the same animal. Overall group effects were assessed using maximum-likelihood likelihood-ratio tests. Pairwise group comparisons were evaluated using two-sided Wald z-tests. Figures were plotted using OriginPro (OriginLab). Statistical analyses were performed using OriginPro for AD-STDM related experiments, GraphPad Prism for Open field, elevated plus maze and fear conditioning experiments, and custom Python scripts for cFos quantification experiments.

## References

1. Radzicki, D., McCann, K. E., Alexander, G. M. & Dudek, S. M. Hippocampal area CA2 activity supports social investigation following an acute social stress. Mol. Psychiatry 30, 2284–2296 (2025).

2. Hung, L. W. et al. Gating of social reward by oxytocin in the ventral tegmental area. Science (1979). 357, 1406–1411 (2017).

3. Green, M. F. & Horan, W. P. Social Cognition in Schizophrenia. 10.1177/0963721410377600 doi:10.1177/0963721410377600.

4. Guo, J. Y., Ragland, J. D. & Carter, C. S. Memory and cognition in schizophrenia. Molecular Psychiatry 2018 24:5 24, 633–642 (2018).

5. Lissek, S. et al. Generalized Anxiety Disorder Is Associated With Overgeneralization of Classically Conditioned Fear. Biol. Psychiatry 75, 909–915 (2014).

6. Hitti, F. L. & Siegelbaum, S. A. The hippocampal CA2 region is essential for social memory. Nature 508, 88–92 (2014).

7. Oliva, A., Fernández-Ruiz, A., Leroy, F. & Siegelbaum, S. A. Hippocampal CA2 sharp-wave ripples reactivate and promote social memory. Nature 587, 264–269 (2020).

8. Meira, T. et al. A hippocampal circuit linking dorsal CA2 to ventral CA1 critical for social memory dynamics. Nat. Commun. 9, 1–14 (2018).

9. Williams Avram, S. K., et al. NMDA Receptor in Vasopressin 1b Neurons Is Not Required for Short-Term Social Memory, Object Memory or Aggression. Front. Behav. Neurosci. 13, 218 (2019).

10. Lein, E. S., Callaway, E. M., Albright, T. D. & Gage, F. H. Redefining the boundaries of the hippocampal CA2 subfield in the mouse using gene expression and 3-dimensional reconstruction. Journal of Comparative Neurology 485, 1–10 (2005).

11. Dudek, S. M., Alexander, G. M. & Farris, S. Rediscovering area CA2: Unique properties and functions. Nature Reviews Neuroscience vol. 17 89–102 Preprint at 10.1038/nrn.2015.22 (2016).

12. Boyle, L. M. et al. The ventral CA2 region of the hippocampus and its differential contributions to social memory and social aggression. Cell Rep. 44, (2025).

13. Fanselow, M. S. & Dong, H. W. Are the Dorsal and Ventral Hippocampus Functionally Distinct Structures? Neuron 65, 7–19 (2010).

14. Leroy, F. et al. A circuit from hippocampal CA2 to lateral septum disinhibits social aggression. Nature 564, 213–218 (2018).

15. Pothuizen, H. H. J., Zhang, W. N., Jongen-Rêlo, A. L., Feldon, J. & Yee, B. K. Dissociation of function between the dorsal and the ventral hippocampus in spatial learning abilities of the rat: A within-subject, within-task comparison of reference and working spatial memory. European Journal of Neuroscience 19, 705–712 (2004).

16. Moser, M. B., Moser, E. I., Forrest, E., Andersen, P. & Morris, R. G. M. Spatial learning with a minislab in the dorsal hippocampus. Proc. Natl. Acad. Sci. U. S. A. 92, 9697–9701 (1995).

17. Ciocchi, S., Passecker, J., Malagon-Vina, H., Mikus, N. & Klausberger, T. Selective information routing by ventral hippocampal CA1 projection neurons. Science (1979). 348, 560–563 (2015).

18. Kjelstrup, K. G. et al. Reduced fear expression after lesions of the ventral hippocampus. Proc. Natl. Acad. Sci. U. S. A. 99, 10825–10830 (2002).

19. Maren, S. & Holt, W. G. Hippocampus and Pavlovian fear conditioning in rats: muscimol infusions into the ventral, but not dorsal, hippocampus impair the acquisition of conditional freezing to an auditory conditional stimulus. Behavioral neuroscience 118, 97–110 (2004).

20. Jimenez, J. C. et al. Contextual fear memory retrieval by correlated ensembles of ventral CA1 neurons. Nat. Commun. 11, 3492 (2020).

21. Kim, W. Bin & Cho, J. H. Encoding of contextual fear memory in hippocampal–amygdala circuit. Nature Communications 2020 11:1 11, 1382-(2020).

22. Kassraian, P. et al. The hippocampal CA2 region discriminates social threat from social safety. Nat. Neurosci. 27, 2193–2206 (2024).

23. Zorab, J. M. et al. Serotonin and neurotensin inputs in the vCA1 dictate opposing social valence. Nature 1–11 (2025) doi:10.1038/s41586-025-08809-2.

24. Suto, N. et al. Neural circuits for valence updating in social memory. Science (1979). 393, (2026).

25. Ryherd, G. L., Bunce, A. L., Edwards, H. A., Baumgartner, N. E. & Lucas, E. K. Sex differences in avoidance behavior and cued threat memory dynamics in mice: Interactions between estrous cycle and genetic background. Horm. Behav. 156, 105439 (2023).

26. Li, L. et al. Social trauma engages lateral septum circuitry to occlude social reward. Nature 613, 696–703 (2022).

27. Li, H. et al. Spatiotemporal Mapping of the Oxytocin Receptor at Single-Cell Resolution in the Postnatally Developing Mouse Brain. Neurosci. Bull. 41, 224 (2024).

28. Zhao, M., Choi, Y. S., Obrietan, K. & Dudek, S. M. Synaptic Plasticity (and the Lack Thereof) in Hippocampal CA2 Neurons. Journal of Neuroscience 27, 12025–12032 (2007).

29. Samadi, M. et al. Mechanisms of mGluR-dependent plasticity in hippocampal area CA2. Hippocampus 33, 730–744 (2023).

30. Evans, P. R. et al. RGS14 Restricts Plasticity in Hippocampal CA2 by Limiting Postsynaptic Calcium Signaling. eNeuro 5, (2018).

31. Lee, S. E. et al. RGS14 is a natural suppressor of both synaptic plasticity in CA2 neurons and hippocampal-based learning and memory. Proc. Natl. Acad. Sci. U. S. A. 107, 16994–16998 (2010).

32. View of c-Fos and neuronal plasticity: the aftermath of Kaczmarek’s theory | Acta Neurobiologiae Experimentalis. Acta Neurobiol Exp 287–298 (2018) doi: 10.21307/ane⍰2018⍰027

33. Koolhaas, J. M. et al. The Resident-intruder Paradigm: A Standardized Test for Aggression, Violence and Social Stress. JoVE (Journal of Visualized Experiments) e4367 (2013) doi:10.3791/4367.

