## Supplemental Figures for "Hippocampal dorsal and ventral CA2 enable social discrimination after acute social defeat"

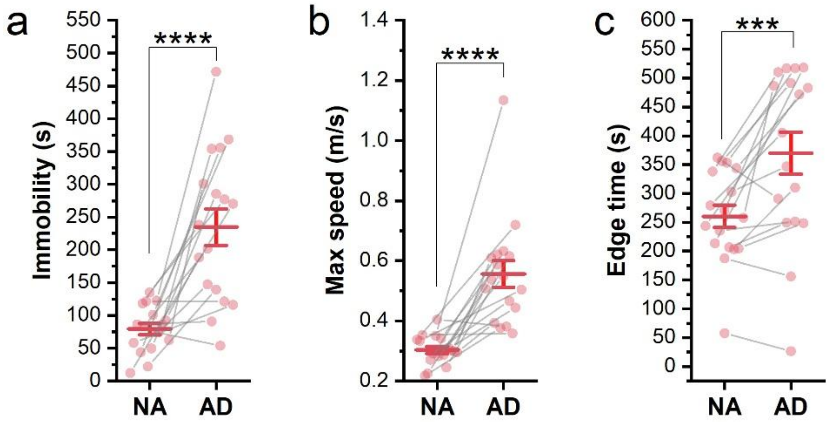


**Supplementary Figure 1: Behavioral differences between NA social interaction and AD on day1.**

**a.** Subject mice spent more time immobile during AD than during NA social interaction. N=17; paired t test, $t = -5.54$, $p < 0.0001$.

**b.** Maximum movement speed was higher during AD than during NA social interaction. N=17; paired t test, $t = -5.69$, $p < 0.0001$.

**c.** Subject mice spent more time in the edge zone of the cage during AD than during NA social interaction. N=17; paired t test, $t = -4.17$, $p = 0.0007$.

***p<0.001, ****p<0.0001

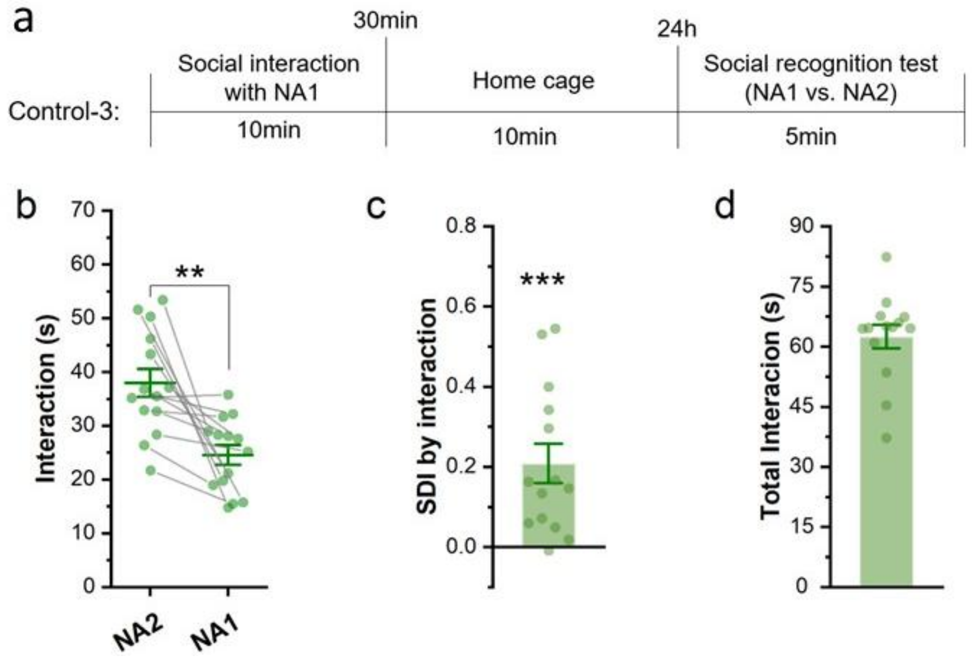


**Supplementary Figure 2: Non-aggressive social memory maintained 24h after the interaction.**
**a.** Schematic of control-3. On day 1, subject mice underwent a 10-min social interaction with a non-aggressor BALB/c mouse (NA1). 24 hours later, the subject mice underwent a 5-min social-recognition test in which the familiar NA1 and a novel NA BALB/c mouse (NA2) were confined in opposite side chambers.
**b.** Subject mice spent more time interacting with the novel NA2 than with the familiar NA1 (paired t test, n = 14, t = -3.93, p = 0.002).
**c.** SDI was higher than 0 for control-3 (one-sample t test vs. 0: t = 4.23, p = 0.0004)
**d.** Total interaction duration.
**p<0.01, ***p<0.001

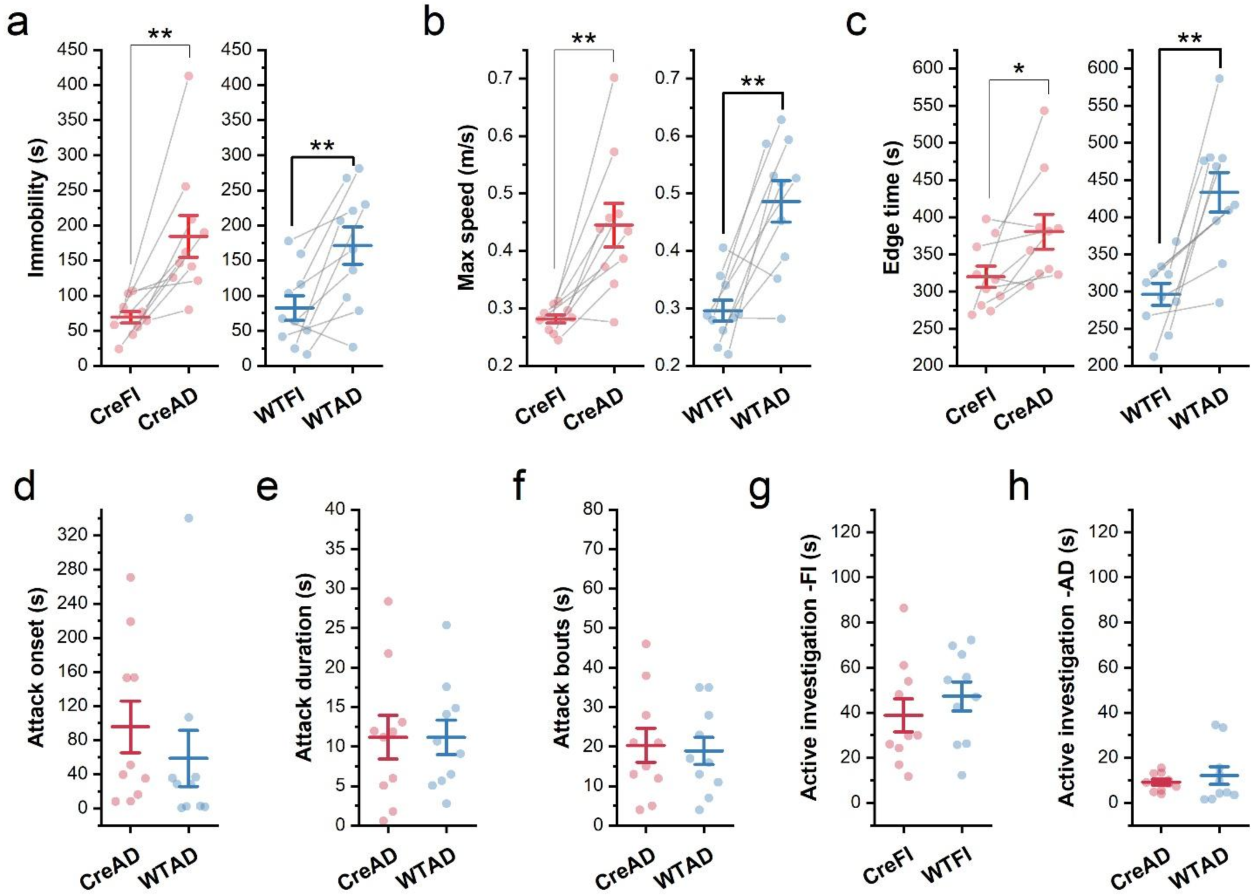


**Supplementary Figure 3. Day 1 behavioral performance during chemogenetic inhibition of Avpr1b-expressing dCA2 neurons.**

**a–c.** Within both the Cre and WT groups, mice displayed longer immobility durations (**a**) (Cre: $t = -3.68$, $p = 0.005$; WT: $t = -3.96$, $p = 0.003$), higher maximum movement speeds (**b**) (Cre: $t = 4.25$, $p = 0.002$; WT: $t = 4.58$, $p = 0.001$), and longer durations in the cage edge zone (**c**) (Cre: $t = 2.31$, $p = 0.046$; WT: $t = 4.699$, $p = 0.001$) during AD than during non-aggressive free interaction (FI).

**d–f.** Attack onset (**d**), total attack duration (**e**), and number of attack bouts (**f**) performed by the aggressor toward the subject mouse during AD did not differ between the Cre and WT groups.

**g–h.** Active investigation towards the stimulus mice did not differ between the Cre and WT groups during either non-aggressive free interaction (FI; **g**) or AD (**h**).

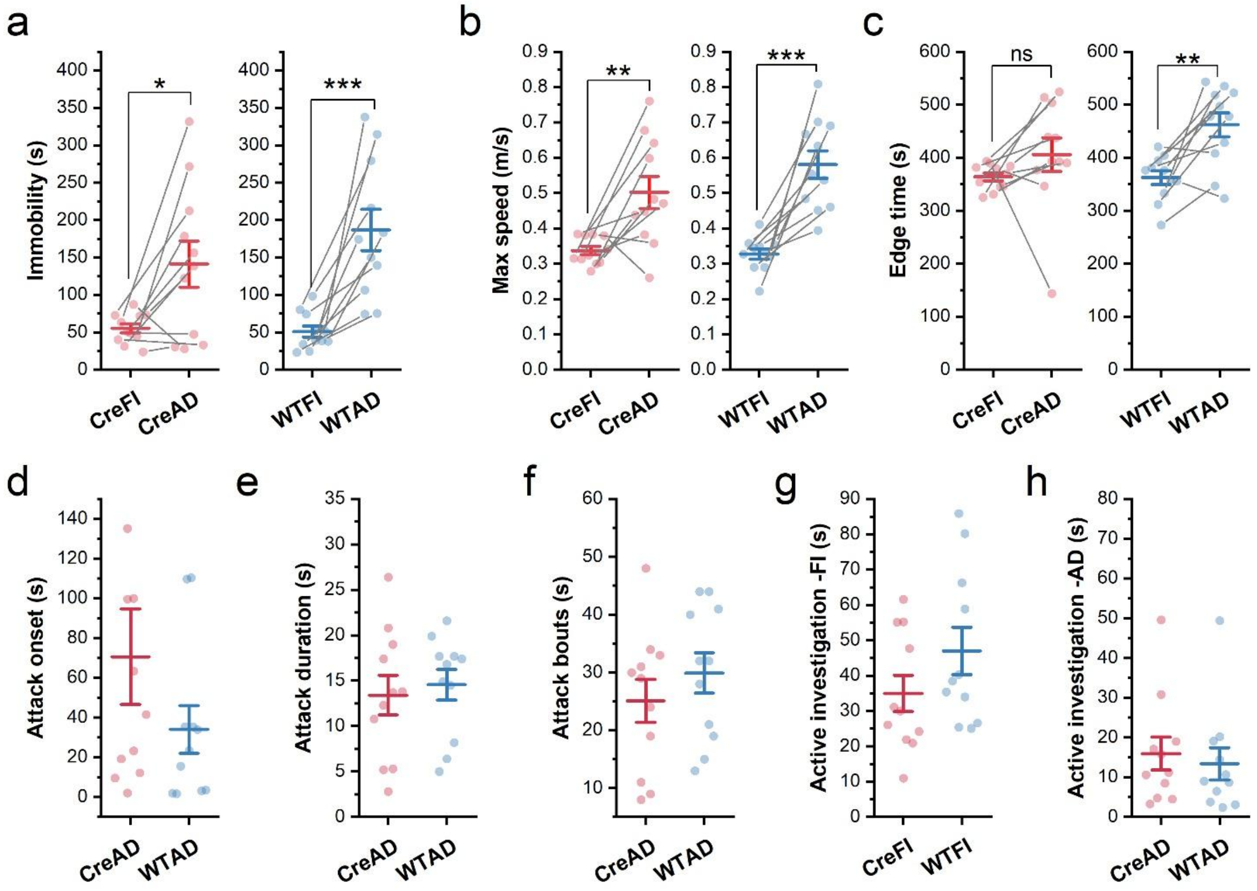


**Supplementary Figure 4. Day 1 behavioral performance in mice subsequently receiving chemogenetic inhibition of Avpr1b-expressing dCA2 neurons during memory recall.**

**a–c.** Within both the Cre and WT groups, mice displayed longer immobility durations (**a**) (Cre: t = -2.81, p = 0.019; WT: t = -5.28, p = 0.0004) and higher maximum movement speeds (**b**) (Cre: t = -3.23, p = 0.008; WT: t = -6.07, p = 0.0001). Edge-zone duration did not differ significantly in Cre mice but was higher during AD than during non-aggressive free interaction (FI) in WT controls (**c**) (Cre: t = -1.2, p = 0.25; WT: t = -4.398, p = 0.0013).

**d–f.** Attack onset (**d**), total attack duration (**e**), and number of attack bouts (**f**) performed by the aggressor toward the subject mouse during AD did not differ between the Cre and WT groups.

**g–h.** Active investigation towards the stimulus mice did not differ between the Cre and WT groups during either non-aggressive free interaction (FI; **g**) or AD (**h**).

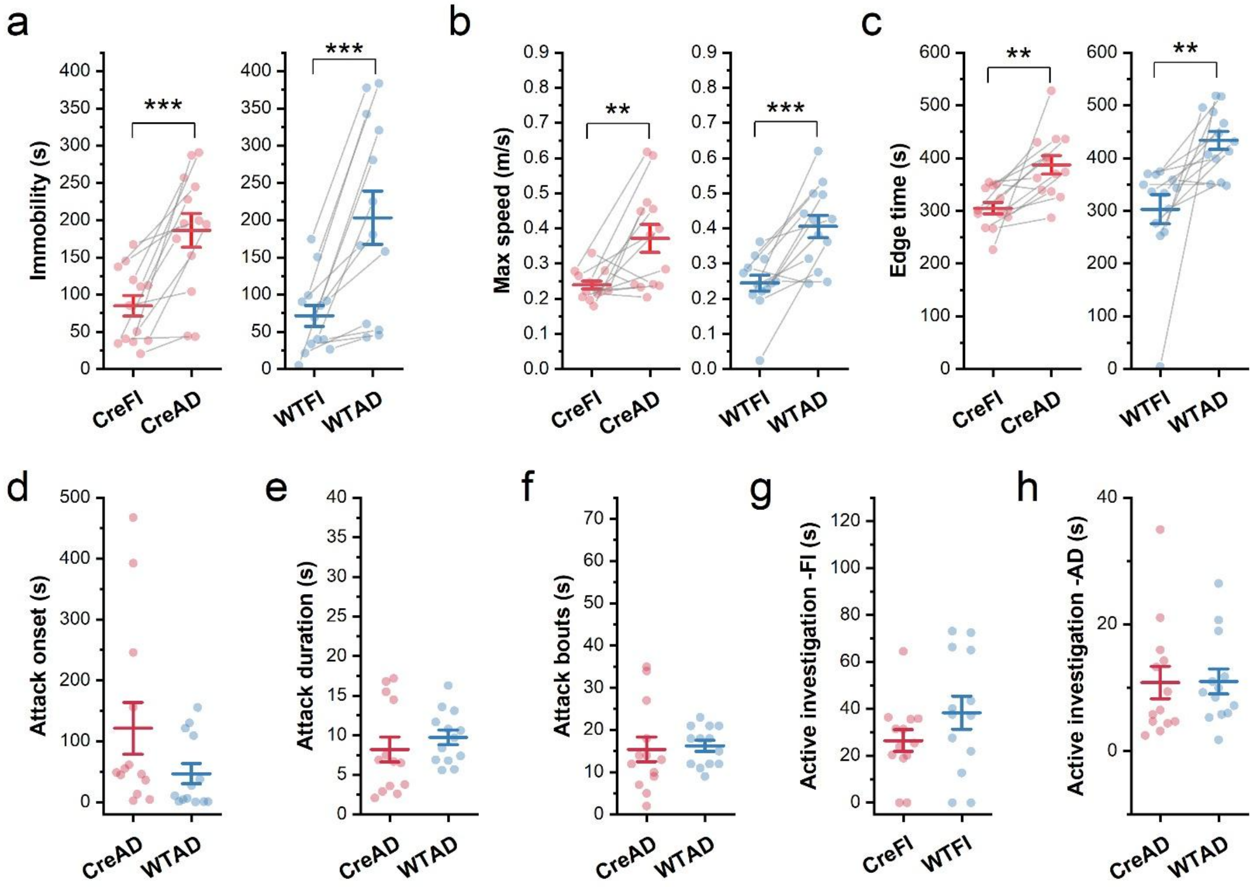


**Supplementary Figure 5. Day 1 behavioral performance during chemogenetic inhibition of Avpr1b-expressing vCA2 neurons.**

**a–c.** Within both the Cre and WT groups, mice displayed longer immobility durations (**a**) (Cre: $t = -4.69$, $p = 0.0005$; WT: $t = -4.55$, $p = 0.0007$), higher maximum movement speeds (**b**) (Cre: $t = -3.18$, $p = 0.008$; WT: $t = -4.44$, $p = 0.0008$), and longer durations in the cage edge zone (**c**) (Cre: $t = -3.76$, $p = 0.003$; WT: $t = -3.39$, $p = 0.005$) during AD than during non-aggressive free interaction (FI).

**d–f.** Attack onset (**d**), total attack duration (**e**), and number of attack bouts (**f**) performed by the aggressor toward the subject mouse during AD did not differ between the Cre and WT groups.

**g–h.** Active investigation towards the stimulus mice did not differ between the Cre and WT groups during either non-aggressive free interaction (FI; **g**) or AD (**h**).

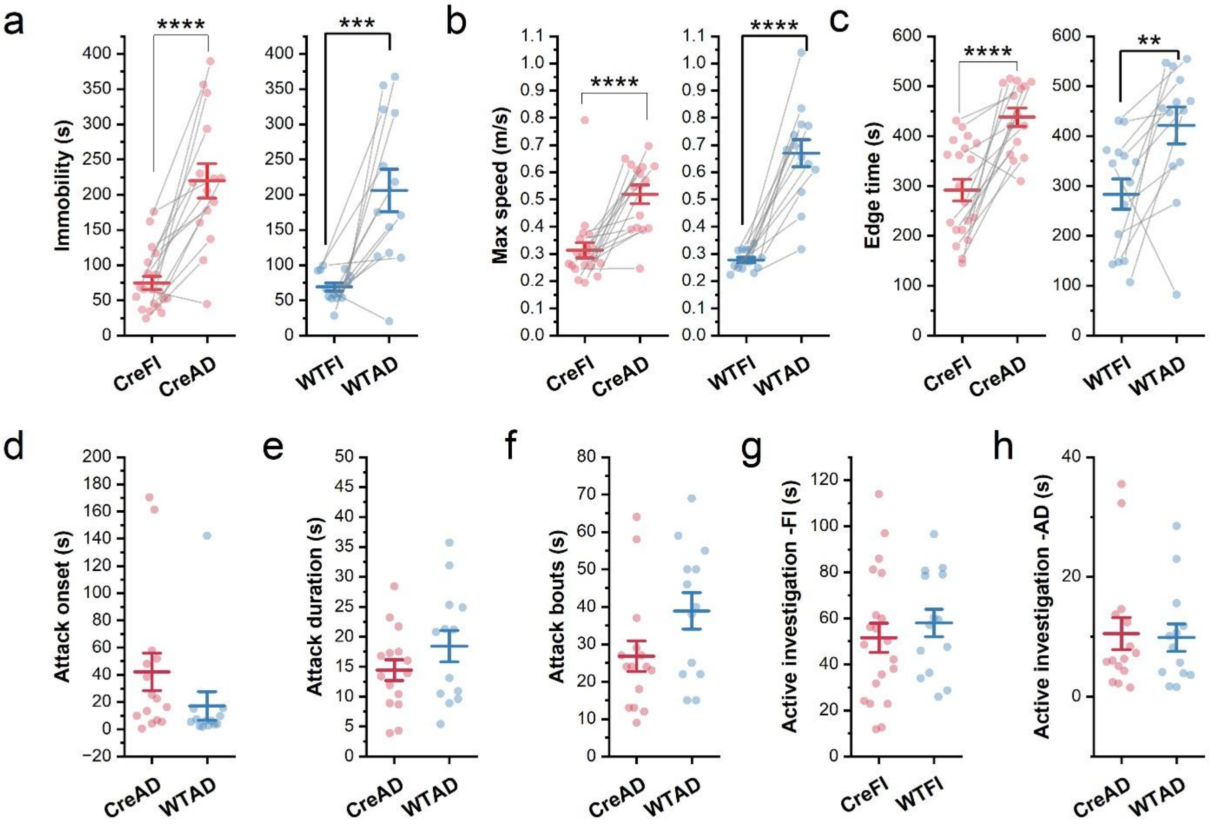


**Supplementary Figure 6. Day 1 behavioral performance in mice received chemogenetic inhibition of Avpr1b-expressing vCA2 neurons during memory recall.**

**a–c.** Within both the Cre and WT groups, mice displayed longer immobility durations (**a**) (Cre: t = -5.44, $p < 0.0001$; WT: t = -4.5, p = 0.0007), higher maximum movement speeds (**b**) (Cre: t = -6.14, $p < 0.0001$; WT: t = -8.58, $p < 0.0001$), and longer durations in the cage edge zone (**c**) (Cre: t = -5.92, $p < 0.0001$; WT: t = -3.11, p = 0.009) during AD than during non-aggressive free interaction (FI).

**d–f.** Attack onset (**d**), total attack duration (**e**), and number of attack bouts (**f**) performed by the aggressor toward the subject mouse during AD did not differ between the Cre and WT groups.

**g–h.** Active investigation towards the stimulus mice did not differ between the Cre and WT groups during either non-aggressive free interaction (FI; **g**) or AD (**h**).

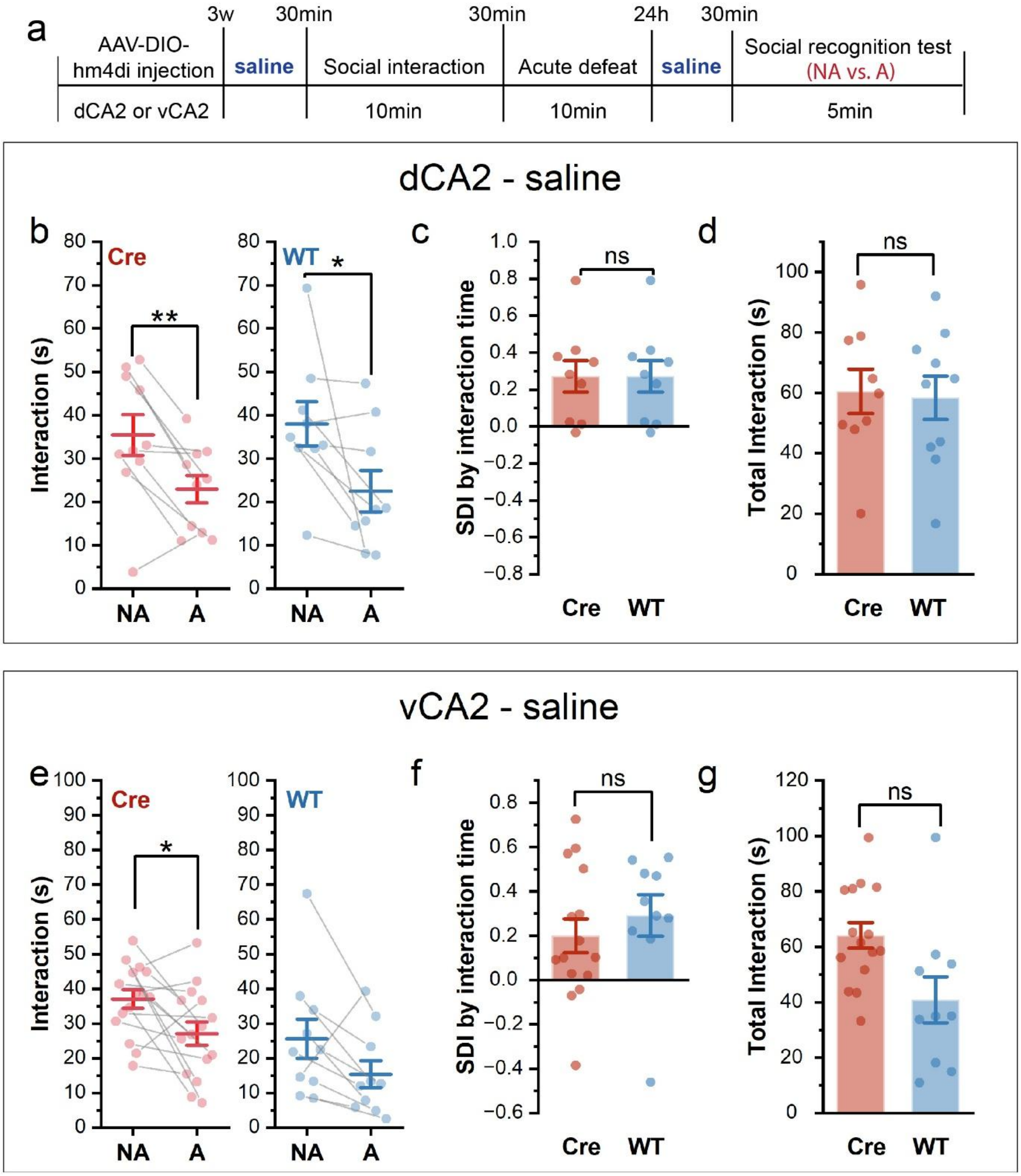
a
3w
30min
30min
24h
30min
AAV-DIO-hm4di injection
saline
Social interaction
Acute defeat
saline
Social recognition test (NA vs. A)
dCA2 or vCA2
10min
10min
5min
dCA2 - saline
b
c
d
Cre
WT
Interaction (s)
NA
A
SDI by interaction time
Total Interaction (s)
ns
vCA2 - saline
e
f
g

**Supplementary Figure 7. Cre and WT group displayed no difference with saline administration in mice received hm4di brain injections.**
**a.** Experimental timeline. AAV-DIO-hM4Di-mCherry was injected into dCA2 or vCA2 of *Avpr1b-Cre*$^{+/-}$ mice and wild-type (WT) controls. Three weeks after viral injection, saline was administered 30 min before both the day 1 learning session and the day 2 recall session. Subject mice then underwent an AD-STDM paradigm.
**b-d. dCA2 saline controls.** Both Cre and WT mice spent more time interacting with the familiar NA than with the A (**b**, n = 9 per group, paired t test, Cre t = 3.56, p = 0.006; WT t = 2.40, p = 0.043). Social discrimination index calculated from the direct interaction time did not differ between the Cre and WT groups (**c**, two-sample t test, t = 0.73, p = 0.48). Total interaction time did not differ between the Cre and WT groups (**d**).
**e-g. vCA2 saline controls.** Cre mice (n = 15) spent more time interacting with the familiar NA than with the A (t = 2.53, p = 0.024), whereas the corresponding comparison in WT mice (n = 10) did not reach statistical significance (t = 2.06, p = 0.07; **e**). Social discrimination index calculated from the direct interaction time did not differ between the Cre and WT groups (**f**, two-sample t test, t = 0.76, p = 0.46). Total interaction time did not differ between the Cre and WT groups (**g**).
ns, not significant; *p < 0.05, **p < 0.01.

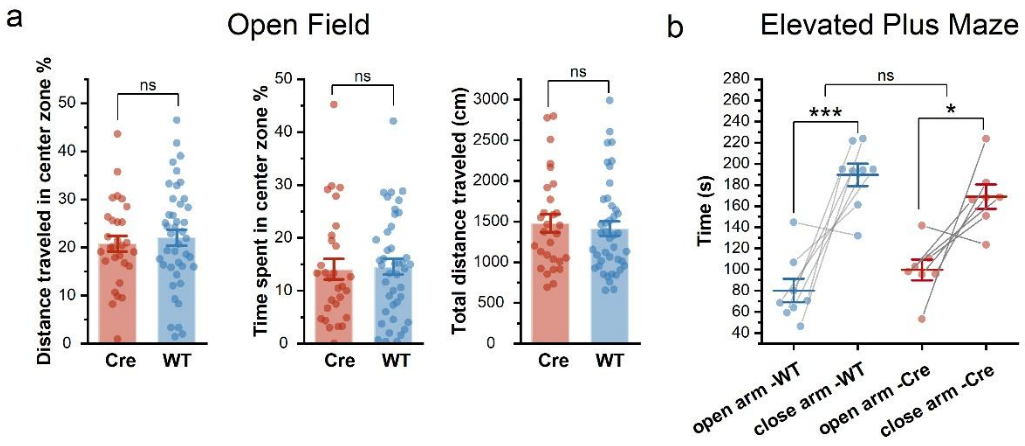


**Supplementary Figure 8: Chemogenetic inhibition of vCA2 Avpr1b neurons does not modulate anxiety-like behavior**

*Avpr1b-Cre*$^{+/-}$ mice were injected with a Cre-dependent hM4Di-mCherry. On the experiment day, mice were injected with CNO. After 30 minutes, mice were run through either a) an open-field test (WT n=42, Cre n=30), or b) an elevated plus maze (WT n=8, Cre n=7) to assess anxiety-like behavior responses.

**a.** No difference was observed in the open field test between *Avpr1b-Cre*$^{-/-}$ and *Cre*$^{+/-}$ mice in terms of distance traveled in the center zone (two-sample unpaired t-test t=0.5126, df=69, p=0.61), time in the center zone (two-sample unpaired t-test t=0.2000, df=69, p=0.84), or total distance traveled (two-sample unpaired t-test t=0.4554, df=69, p=0.65).

**b.** No difference was observed between groups for time spent in the open or closed arm of the elevated plus maze (Two-way repeated measures ANOVA for Genotype x Time in Arm F(1,13)=1.715, p=0.21), and both groups showed significant preference for the closed arms over the open arms (Two-way repeated measures ANOVA for Time in Arm F(1,13)=34.11, p<0.0001, Šídák's multiple comparisons test WT p=0.0003, Cre p=0.017).

ns, not significant; *p < 0.05, ***p < 0.001.

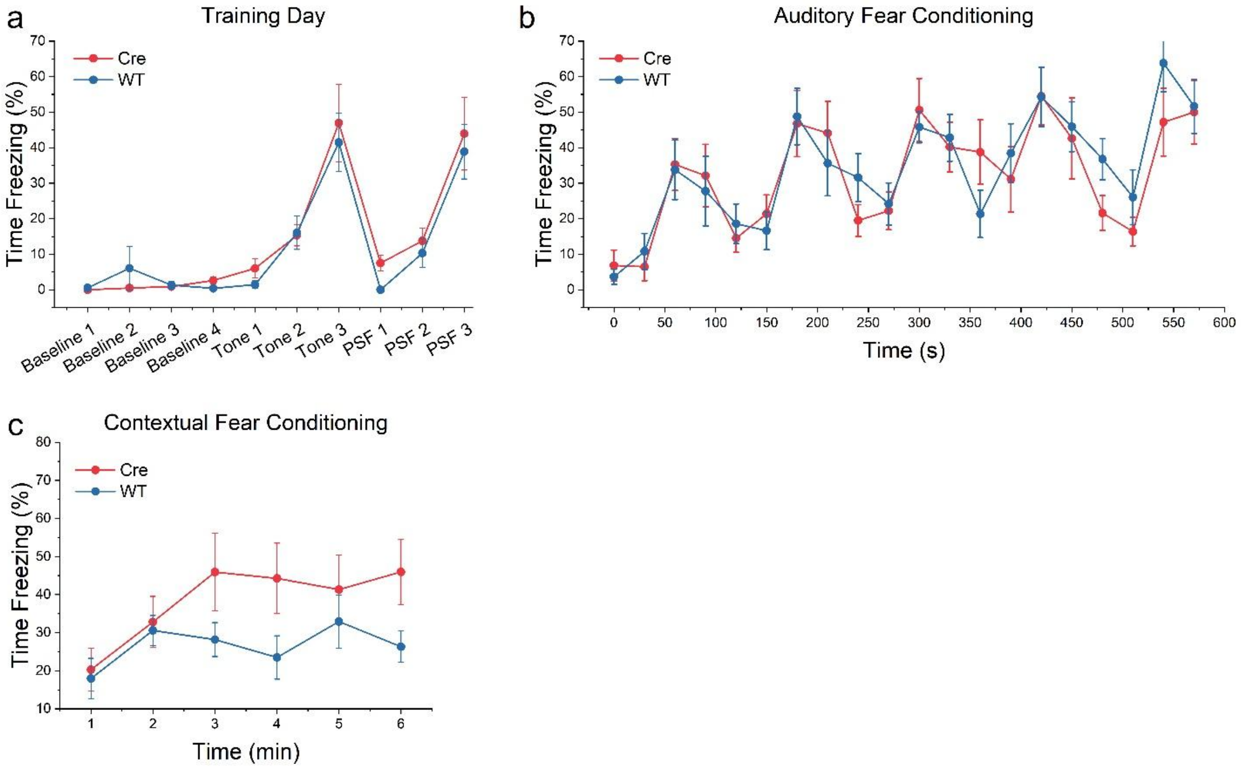


**Supplementary Figure 9: Chemogenetic inhibition of vCA2 Avpr1b neurons does not alter auditory or contextual fear conditioning**

**a.** *Avpr1b-Cre*$^{+/-}$ (n=9) and *Cre*$^{-/-}$ WT (n=8) mice were injected with a Cre-dependent hM4Di-mCherry. On the experimental day, mice were injected 30 minutes prior to testing with 5mg/kg CNO intra-peritoneally. On training day, mice were exposed to tone and shock pairings in context A. No difference was observed in time spent freezing across conditions between WT and Cre animals. Two-way repeated-measures ANOVA Genotype x Period $F(9,135)=0.4298$, $p=0.92$

**b.** On the first test day (24 hours post-training), mice were exposed to context B, and the time spent freezing following multiple tones was quantified. No significant difference in freezing was observed between groups. Two-way repeated-measures ANOVA Genotype x Time $F(19,285)=1.036$, $p=0.42$

**c.** On the second test day (48 hours post-training), mice were exposed to the same context A, and the time spent freezing across time was quantified. No significant difference in percent time freezing was observed across time between groups. Two-way repeated-measures ANOVA Genotype x Time $F(5,75)=1.484$, $p=0.21$

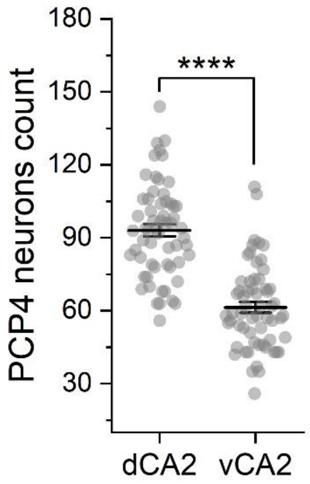


**Supplementary Figure 10: Hippocampal dCA2 contains more PCP4-expression neurons than vCA2.**

Quantification of PCP4+ neurons in the anatomically identified CA2 regions. (n = 12 mice and 60 slices, with five slices per mouse. Unpaired t test: dCA2 vs. vCA2, t = 9.51, p < 0.0001)

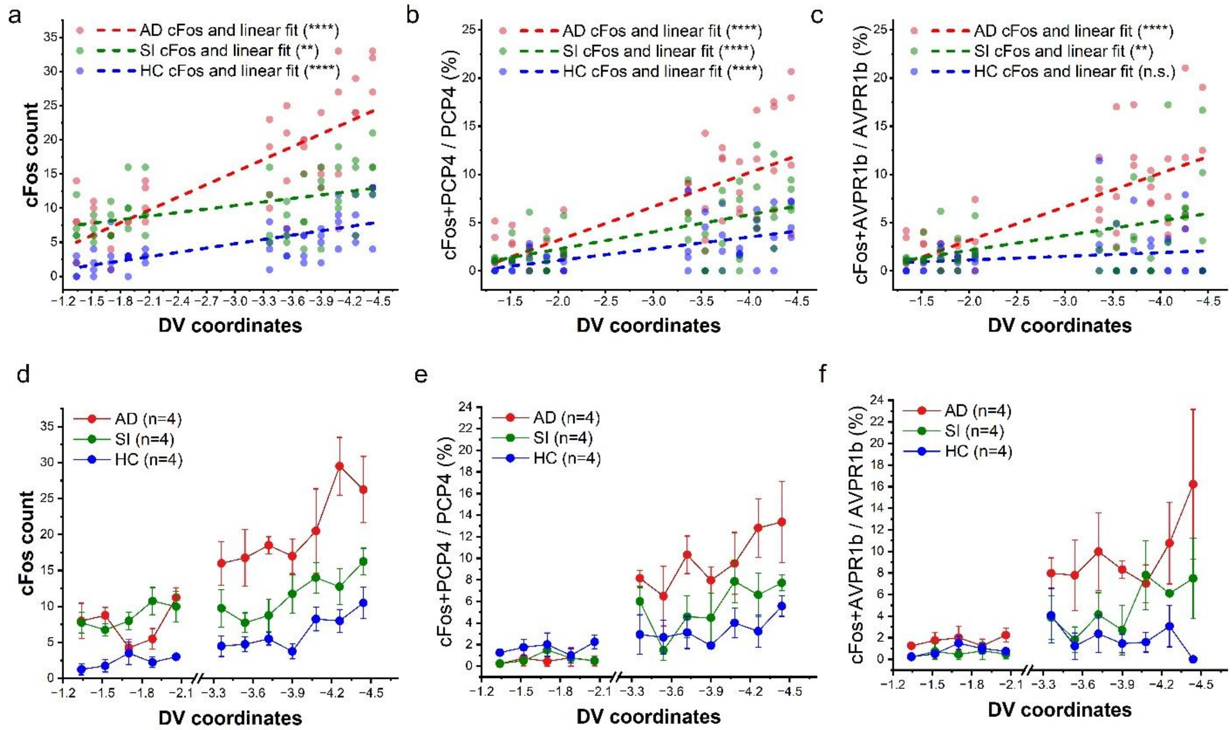


**Supplementary Figure 11: cFos expression varies as a function of dorsoventral coordinates.**
**a–c.** cFos expression was positively correlated with position along the dorsoventral axis of CA2 in NeuroTrace+ neurons (**a**), PCP4-expressing neurons (**b**), and Avpr1b-expressing neurons except the HC group (**c**). *n* = 4 slices per group at each coordinate. Slices were collected from the same four mice.
**d–f.** Analysis across dCA2 and vCA2 revealed a significant overall effect of group: cFos expression was higher in the AD group than in the SI group in vCA2, while no such difference was detected in dCA2. Dorsal (−1.34 to −2.06 mm) and ventral (−3.54 to −4.44 mm) CA2 were analyzed separately using two-way mixed-design repeated-measures ANOVA with Holm test. (**d**), total cFos counts (dCA2: $F(2, 9) = 21.952$, $p = 0.000345$; HC vs. SI, $p = 0.000475$; HC vs. AD, $p = 0.00126$; SI vs. AD, $p = 0.307$; vCA2: $F(2, 9) = 14.041$, $p = 0.00171$; HC vs. SI, $p = 0.103$; HC vs. AD, $p = 0.00165$; SI vs. AD, $p = 0.0156$); (**e**), PCP4-normalized cFos (dCA2: $F(2, 9) = 19.536$, $p = 0.000532$; HC vs. SI, $p = 0.00230$; HC vs. AD, $p = 0.000664$; SI vs. AD, $p = 0.244$; vCA2: $F(2, 9) = 10.682$, $p = 0.00420$; HC vs. SI, $p = 0.202$; HC vs. AD, $p = 0.00441$; SI vs. AD, $p = 0.0241$); (**f**), Avpr1b-normalized cFos (dCA2: $F(2, 9) = 48.053$, $p < 0.0001$; HC vs. SI, $p < 0.0001$; HC vs. AD, $p < 0.0001$; SI vs. AD, $p = 0.0979$; vCA2: $F(2, 9) = 16.341$, $p = 0.00101$; HC vs. SI, $p = 0.0469$; HC vs. AD, $p = 0.000902$; SI vs. AD, $p = 0.0162$).
$^{**}p<0.01$, $^{***}p<0.001$, $^{****}p<0.0001$
